# Spatial (Mis)match Between EEG and fMRI Signal Patterns Revealed by Spatio-Spectral Source-Space EEG Decomposition

**DOI:** 10.1101/2024.07.26.605087

**Authors:** Stanislav Jiricek, Vlastimil Koudelka, Dante Mantini, Radek Marecek, Jaroslav Hlinka

**Author notes:** **Correspondence** Jaroslav Hlinka, Department of Complex Systems, Institute of Computer Science of the Czech Academy of Sciences, Czech Republic.

## Abstract

In this work, we aimed to directly compare and integrate EEG whole-brain patterns of neural dynamics with concurrently measured fMRI BOLD data. For that purpose, we set out to derive EEG patterns based on a spatio-spectral decomposition of band-limited EEG power in the source-reconstructed space. On a large data set of 72 subjects resting-state hdEEG-fMRI we showed that the proposed approach is reliable both in terms of the extracted patterns as well as their spatial BOLD signatures. The five most robust EEG spatio-spectral patterns include, but go beyond, the well-known occipital alpha power dynamics. The EEG spatial-spectral patterns show relatively weak, yet statistically significant spatial similarity to their fMRI BOLD signatures, particularly the patterns that show stronger temporal synchronization with BOLD. However, we observed an insignificant relation between the temporal synchronization and spatial overlap of the EEG spatio-spectral patterns and the classical fMRI BOLD resting state networks (as obtained by independent component analysis). This provides evidence that both EEG (frequency-specific) power and BOLD signal capture reproducible spatiotemporal patterns of neural dynamics. Rather than being mutually redundant, these are only partially overlapping, carrying to a large extent complementary information concerning the underlying low-frequency dynamics. Finally, we report and interpret the most stable source space EEG-fMRI patterns, along with the corresponding EEG electrode space patterns better known from the literature.

## 1 INTRODUCTION

To find a reliable link between electroencephalography (EEG) and functional magnetic resonance imaging (fMRI) data measured during resting brain seems to be similarly difficult as to fulfill the “to rest” instruction for the volunteering subject. Yet, there is variety of strong reasons for attempting to fuse the two modalities. First, finding a relation may help in understanding the origin and character of each type of signal. Second, by utilizing simultaneous EEG-fMRI measurement, we could suppress weaknesses of both modalities when using them separately and thus study the healthy brain as well as brain disorders more accurately. The third motivation is the search for aspects of EEG signal that would reproduce the well-described features of the BOLD signal - the resting-state networks (RSNs) dynamics. Discovering such relations could in principle in future allow replacing in more scenarios the costly fMRI measurements by the much more affordable EEG experiments. This could ultimately provide more affordable diagnostics and treating methods (Vega, Michel, Saxena, White, & Valdes-Sosa 2022), increase ecological validity of experiments as well as enable wider range of experiment designs.

Since (Logothetis, Pauls, Augath, Trinath, & Oeltermann 2001) experimentally showed a delayed hemodynamic response of the Blood-oxygen-level-dependent (BOLD) fMRI signal to the measured local field potential (LFP) signal envelope, questions concerning the details of the relation of electrophysiological signal and fMRI have been around and are still far from satisfactorily answered, particularly when one moves outside of the well controlled situation of a controlled task experiment and localized intracortical EEG recording. The standard noninvasive EEG signal can be naturally described in terms of three domains: temporal, spatial, and frequency domain. Initial resting-state studies aimed to find the BOLD correlate of the most prominent resting-state EEG feature, i.e., the occipital alpha activity (Berger 1929). In those studies, the spatial domain was mostly reduced by selecting the occipital subset of electrodes, and similarly the frequency domain was limited to the alpha frequency band by utilizing a band-limited power (BLP) regressor for BOLD signal prediction (Feige et al. 2005; Goldman, Stern, Engel Jr, & Cohen 2002; Moosmann et al. 2003). Other studies varied in terms of the used electrode subsets (Laufs, Kleinschmidt, et al. 2003) or frequency bands (Laufs, Krakow, et al. 2003), and also the potential use of multiple frequency specific regressors in the same statistical general linear model (GLM) (de Munck, Gonçalves, Mammoliti, Heethaar, & Da Silva 2009; Tyvaert, LeVan, Grova, Dubeau, & Gotman 2008). Despite a very similar experimental design those initial studies generally found (one of) two distinct spatial BOLD fMRI correlation patterns of the alpha regressor. Namely, a widespread bilateral fronto-parietal correlation pattern was reported by (Laufs, Kleinschmidt, et al. 2003), while (Goldman et al. 2002; Moosmann et al. 2003) observed an occipito-parietal pattern. Thus, while both patterns showed negative relation between the EEG alpha BLP regressor and the BOLD signal (which was in line with the intuitive interpretation of alpha as the iddling rhythm corresponding to low metabolism requirements), the localization of these anticorrelations differed substantially between the studies, creating a confusion concerning which brain areas were actually most involved in this iddling. Later, by reanalyzing their data, Laufs et al. (2006) showed that both these alpha correlates can be observed within the same data set when considering different subset of subjects. Similarly, Gonçalves et al. (2006) showed that the alpha correlate shows a high level of the inter-as well as intra-subject variability.

Other studies moved the focus of the field to other frequency bands than alpha: Scheeringa, Petersson, Kleinschmidt, Jensen, and Bastiaansen (2012) showed that frontal theta rhythm power fluctuation positively correlates with hubs of the default mode network (DMN). Further, (Mantini, Perrucci, Del Gratta, Romani, & Corbetta 2007) found that BLP time series at standard EEG frequency bands taken as an average signal across all electrodes correlates with time series of multiple BOLD independent components (ICs), i.e., resting state networks (RSNs). This as well as previous findings suggest that the dynamics of RSNs might be reflected in EEG as a combination of multiple frequency and spatial patterns. The first EEG-fMRI data-driven method that did not put any a priori assumptions on spatial, temporal or frequency domain was introduced in (Miwakeichi et al. 2004) and subsequently in simultaneous EEG-fMRI study (Martınez-Montes, Valdés-Sosa, Miwakeichi, Goldman, & Cohen 2004) where the three-dimensional spatio-temporal-frequency array was factorized by the parallel factor analysis (PARAFAC) (Harshman & Lundy 1994) to obtain spatio-temporal-frequency components. In correspondence with (Goldman et al. 2002), the authors found a single alpha band occipital component that had significant temporal correspondence with the BOLD signal in the occipital lobe. The authors also reported a frontal theta component in correspondence with (Scheeringa et al. 2012), which did not have a significant correlation with the BOLD signal. Similar analysis was also performed in (Mareček et al. 2017; Marecek et al. 2016), where several components representing brain rhythms as well as artifacts were identified. Even though those trilinear decomposition methods represent true data-driven approach for identifying EEG spatio-temporal-frequency patterns, they might fail in identifying spatially and frequency-wise distinct BLP fluctuations, because each component in this model has only one spatial signature for all frequency bands and vice versa.

Therefore, it seems more convenient to utilize a model that allows different frequency bands (or combinations thereof) to have different spatial signatures within one component. One of such methods, called spatio-spectral decomposition (Bridwell, Wu, Eichele, & Calhoun 2013) is based on a concatenation of spatial and frequency domain. A matrix decomposition, such as independent component analysis (ICA) (Hyvärinen & Oja 2000), is subsequently applied to this joint spatio-spectral dimension. Bridwell et al. (2013) introduced this concept for simultaneous EEG-fMRI data and linked the obtained components to the BOLD RSNs. Bridwell, Rachakonda, Silva, Pearlson, and Calhoun (2018) further validated several such blind source separation algorithms (BSS) on realistic as well as simulated data. Labounek et al. (2018) reported robust spatio-spectral patterns (SSPs) across three data sets of different paradigms and later (Labounek et al. 2019) a temporal relation of SSPs to the BOLD signal in various hubs of BOLD RSNs. Furthermore, significant relation of several SSPs to the stimuli types of different paradigms was also found (Labounek et al. 2021). A potential drawback of this method is, that the spatial domain is defined by the electrode space which makes the spatial signatures difficult to interpret or to directly compare with fMRI BOLD maps. Readers will notice that here we review only studies within the so-called EEG-informed fMRI integration approach Abreu, Leal, and Figueiredo (2018) as well as studies that derive mostly BLP features from EEG data to predict the BOLD signal. For a broader overview of integration approaches and EEG feature extraction methods, readers are encouraged to refer to comprehensive review articles by (Abreu et al. 2018; Jorge, Van der Zwaag, & Figueiredo 2014; Murta, Leite, Carmichael, Figueiredo, & Lemieux 2015).

Recently, there is an effort to make the high-density EEG (hdEEG) a true neuroimaging tool (as compared with the standard low-density EEG, where the low tens of recording electrodes hardly provide an *image* of brain activity in the computer science sense, and methods applied correspond much more to its time series character). In a recent resting-state study, (Liu, Farahibozorg, Porcaro, Wenderoth, & Mantini 2017) obtained a full set of RSNs from precisely source-localized BLP ICA decomposition similar to the one performed typically on the BOLD signal. In (Liu, Ganzetti, Wenderoth, & Mantini 2018), the authors demonstrated the importance of precisely performed source localization steps, such as precise electrode localization (Marino, Liu, Brem, Wenderoth, & Mantini 2016), multi-layer individual head model to define a reliable forward model (Taberna, Samogin, & Mantini 2021), or the electrode coverage density. In (Marino, Arcara, Porcaro, & Mantini 2019), the authors recently obtained EEG source-reconstructed DMN spatially and temporally related to the BOLD-derived DMN from simultaneous resting-state EEG-fMRI recordings. On the other side, different analysis but with the same goal to obtain DMN characteristics from EEG was shown in (Prestel, Steinfath, Tremmel, Stark, & Ott 2018), where the authors point to the fact that the component having high temporal correspondence with the DMN was associated with eye movement artifacts.

Beside the above mentioned studies, there are other recent studies investigating the link between resting-state EEG and fMRI in the EEG source-reconstructed space. In (Sockeel, Schwartz, Pélégrini-Issac, & Benali 2016), the authors concatenated temporal and frequency domain to perform a matrix decomposition to obtain a separate temporal dynamics for each of 5 frequency bands. They reported a correspondence between EEG and fMRI RSNs, but also substantial mismatches. Similar analysis was also performed in (Li et al. 2018). In (Abreu, Simões, & Castelo-Branco 2020), despite having a data set with only 10 subjects, the authors report a high spatial correspondence between the EEG and fMRI RSNs and also a significant match between the EEG a fMRI dynamical functional connectivity (dFC) states. Another study (Yuan et al. 2016) aimed to obtain the full set of EEG RSNs and compare them spatially and temporally with BOLD RSNs. They found a spatial correspondence but in the same time very low temporal correspondece between EEG and fMRI RSNs. In (Meyer, Janssen, Van Oort, Beckmann, & Barth 2013), the authors claim that the correlation between EEG BLPs and BOLD RSNs is unstable over time.

The goal of this study is to merge literature streams of spatio-spectral decomposition and reliable source localization techniques and introduce a spatio-spectral decomposition in the source-reconstructed space that will benefit from the data-driven nature of the spatio-spectral decomposition and cortical sources spatial domain and allows us to directly test the relationship between those modalities in a way that was not possible before. On a large simultaneous EEG-fMRI data set, we investigate the stability of decomposition in both EEG source and electrode spaces as well as how the EEG patterns relate to BOLD activation patterns and BOLD RSNs, which are all questions of a hot debate in the current literature.

Throughout this manuscript, we emphasize the use of robust evaluation methods that provide an unbiased view of intermodal relationships. Specifically, we begin by examining the stability of EEG decomposition itself (subsection 2.6) to ensure the algorithmic stability of the EEG decomposition. Subsequently, we present a biased version of EEG feature reproducibility testing in subsection 2.9.1 and argue for more sophisticated evaluation approaches that consider not only EEG patterns but also their link to fMRI BOLD data. To address this, we introduce EEG-fMRI integration reproducibility analysis (subsection 2.9.2), assessing the stability of derived EEG patterns and their association with fMRI data. After that, we identify the most stable EEG-fMRI patterns (subsection 2.10). Finally, employing source localization methods, we directly test three hypotheses regarding spatio-temporal relationships between EEG and BOLD data: 1) Are EEG patterns colocalized with their corresponding fMRI BOLD activation maps? (subsection 2.11.1); 2) Do EEG patterns explaining a greater portion of BOLD data variability exhibit higher colocalization? (subsection 2.11.2); 3) Are EEG patterns spatio-temporally associated with BOLD resting-state networks? (subsection 2.11.3). We believe that addressing these questions is essential for developing a more reliable understanding of the EEG/fMRI relationship.

## 2 METHODS

### 2.1 Participants and experimental design

For the purpose of this study, we utilized two data sets. The first and main data set comprises simultaneous EEG-fMRI recordings as well as several types of structural images necessary for constructing individual models of the brain. The second data set consists of out-of-scanner EEG recordings without functional MRI data and this data set serves for illustrating its robustness with respect to change of dataset, preprocessing, and absence of MRI artifacts. Both are described in the following text.

#### 2.1.1 EEG-fMRI data set

We utilized a data set of a single study acquired during a 4 year period in the Central European Institute of Technology (CEITEC) in Brno, Czech Republic. We analyzed 72 healthy participants (mean age: 31.4, range: 18.3 – 50.6, 36 males and 36 females) having consistent data acquisition parameters. All participants underwent a 20 minutes eyes-closed resting-state EEG-fMRI recording session with instructions to lay still, not to fall asleep, and not to think about anything in particular. The study was approved by the ethical committee of Masaryk University and was conducted in accordance with the Declaration of Helsinki. All participants provided written informed consent to participate in the study.

#### 2.1.2 Out-of-scanner EEG data set

To relate the EEG patterns extracted from the combined EEG-fMRI dataset to the patterns obtainable when applying common processing steps to out-of scanner EEG, we utilized the following out-of-scanner EEG data set. The EEG data from healthy subjects were acquired in the National Institute of Mental Health (NIMH, Klecany, Czech Republic) in frame of the multimodal prospective database of first episodes of psychotic illness project. The data set consists of 50 healthy participants(mean age: 29.9, range: 19.1 – 43.0, 21 males and 29 females). Each participant was subjected to a 15-minute resting state paradigm (first 5 minutes eyes open, last 10 minutes eyes closed condition). The 10 minutes of eyes closed resting state recording were preprocessed and analyzed. The study was approved by the local ethical committee of the NIMH and was conducted in accordance with the Declaration of Helsinki. All participants provided written informed consent to participate in the study. The differences in acquisition parameters and processing steps between this out-of-scanner data set and the previously mentioned EEG-fMRI data set are shortly discussed at the end of each of the following subsections. Indeed, in both research and clinical practice, the same acquisition and processing steps can’t be typically guaranteed, and some differences are thus to be expected in most subsequent studies aiming to apply the proposed methodology for EEG SSP extraction to datasets from other EEG experiments. Therefore, this out-of-scanner dataset is not intended for testing replicability under optimal conditions, but rather to serve as an illustrative example, elucidating the extent to which the results observed in the EEG-fMRI data set are robust with respect to applying it to another dataset with established standard, yet slightly different processing.

### 2.2 Data acquisition

The magnetic resonance imaging was performed using a 3T Siemens Prisma magnetic resonance scanner with a 64-channel RF receiving head coil (Siemens Healthineers, Erlangen, Germany). The functional magnetic resonance imaging was performed with a multiband multi-echo 2D EPI sequence with the following parameter settings: Multiband factor: 6; Number of echos: 3; PAT factor: 2; TR = 650 ms; TE = 14.60/33.56/52.52 ms; 48 axial slices with 3 mm slice thickness; slice: 64×64 matrix, 194×194 mm; number of volumes: 1840; FA = 30°. Two structural MRI images were acquired. The first one was obtained without an electrode net with the T1 MPRAGE sequence and the following parameters setting: TR = 2300 ms; TE = 2.34 ms; TI = 900 ms; 240 sagital slices with 1 mm slice thickness; slice: 260×256 matrix, 260×256 mm; FA = 8°; PAT factor: 2. The second structural MRI image with T1 MPRAGE sequence was acquired with the electrode net put on for precise electrode localization and the parameter setting: TR = 2300 ms; TE = 2.33 ms; TI = 900 ms; 240 sagital slices with 1 mm slice thickness; slice: 224×224 matrix, 224×224 mm; FA = 8°; PAT factor: 7. The EEG data with sampling frequency 1000 Hz were recorded with an EGI Hydrocell MR-compatible 256-channel high-density electrode net plugged in the EGI GES 400 signal amplifier (Electrical Geodesics, Inc., Eugene, Oregon, United States). To obtain the ECG signal, one additional channel was recorded. Breathing belt was also attached to the participant’s chest to record breathing cycles.

The out-of-scanner EEG data acquisition was performed with the very same recording set up with the only difference of having the non MR-Compatible version of the 256-channel high-density electrode net. The individual structural T1 MRI MPRAGE image was also provided with the following acquisition parameters: voxel size of 1×1×1 mm, 224 sagittal slices, TE = 4.63 ms, TR = 2300 ms, TI = 900 ms, FA = 10°, TA = 5:30, FOV = 256 mm.

### 2.3 EEG data preprocessing

The raw EEG data were preprocessed by a fully-automated pipeline introduced in (Liu et al. 2017 2018; Marino et al. 2019). The preprocessing pipeline utilizes built-in and in-house MATLAB (MathWorks, Natick, MA, United States) functions as well as SPM (Penny, Friston, Ashburner, Kiebel, & Nichols 2011), Fieldtrip (Oostenveld, Fries, Maris, & Schoffelen 2011), and EEGLAB (Delorme & Makeig 2004) toolboxes. The preprocessing steps are summarised in the following paragraph.

The first step was gradient artifact removal by the FMRI Artifact Slice Template Removal (FASTR) method (Niazy, Beckmann, Iannetti, Brady, & Smith 2005) in EEGLAB followed by a ballistocardiogram artifact removal utilizing adaptive optimal basis set method introduced in (Marino et al. 2018). The bad channels were identified based on low correlation with all other channels in wide frequency band (1 - 80 Hz) and the variance in EEG non-physiological frequency band 200 - 250 Hz. The latter metric serves as a noise variance indicator. In case when at least one criterion marked an outlier in distribution across channels, the channel time course was subsequently interpolated by the time courses of neighboring channels based on the weighted average (electrode distances) scheme implemented in Fieldtrip (Oostenveld et al. 2011). Subsequently, EEG data were filtered in the 1 - 80 Hz frequency band, and ICA was applied to remove movement and other biological artifacts including electrooculographic (EOG) and electromyographic (EMG) artifacts from the EEG recordings. For that purpose FastICA (Hyvärinen & Oja 2000) algorithm based on a deflation approach and hyperbolic tangent as the contrast function was applied and artifactual components were detected based on three parameters, namely the correlation values between ICs time course and reference EOG and EMG signals, the similarity of ICs power spectrum with a 1/f function, and kurtosis of ICs timecourse (Mantini, Franciotti, Romani, & Pizzella 2008). The artifact-suppressed EEG data were subsequently filtered into several frequency bands: Delta (*δ*, 1 - 4 Hz), Theta (*0*, 4 - 8 Hz), Alpha (*α*, 8 - 12 Hz), low Beta (*/J*_1_, 12 - 15 Hz), middle Beta (*/J*_2_, 15 - 18 Hz), high Beta (*/J*_3_, 18 - 30 Hz), and Gamma (*γ*, 30 - 44 Hz). Although the EEG data were comprehensively preprocessed, we decided to exclude all cheek electrodes and the two lowest layers of the neck electrodes from further analysis since there is a concern in current literature that sensors placed at those electrode sites contain disproportionally more artifacts (Vorderwülbecke et al. 2020). Therefore, for all further analyses we utilized 195 out of 257 electrodes. As the last step of the preprocessing, we rereferenced the EEG data to average reference.

The out-of-scanner EEG data were preprocessed manually by the expert with the following steps: EEG data were pre-processed in BESA v. 7.0 software (MEGIS, Munich, Germany) by removing noisy epochs followed by a semi-automated ECG and eye movement-related artifact correction by the signal-space projection method (Uusitalo & Ilmoniemi 1997).

### 2.4 MRI data preprocessing

The functional MRI data were preprocessed in MATLAB utilizing a combination of SPM12 (Penny et al. 2011) pipelines and in-house scripts. At first, the fMRI volumes were spatially realigned, followed by a fusion of 3 echos by a weighted averaging based on a temporal signal-noise-ratio (tSNR). To regress out cardiac and breathing artifactual signals from the BOLD signal, the RETROICOR method (Glover, Li, & Ress 2000) informed by the ECG and breathing signals was employed. Then followed a corregistration of the structural MRI to the average functional image and a structural MRI was normalized to MNI space based on an image segmentation to grey and white matter. Functional data were subsequently transformed to MNI space utilizing the combined transformation matrices from the previous step. All volumes were also resampled into 3×3×mm isotropic voxels. Individual fMRI volumes were then spatially smoothed by a Gaussian filter (FWHM 5 mm). To further mitigate the risk of spurious correlations that can cause common artifacts appearing over the entire volume and having a non-physiological nature, signal from the grey matter was orthogonalized to the following proxies of contributing artifactual signals: a bank of sinusoidal signals with low frequencies (periods slower than 128 s), the first PCA component from the voxel signals of white matter mask, the first PCA component from the voxel signals of CSF mask, set of 24 rotation and translation parameters of estimated head motion, namely the estimated motion parameters themselves, their first differences, squares, and squared first differences. Finally, we applied a low-pass filter to the BOLD data with cut-off frequency 0.09 Hz, which is a typical filtering step for resting-state connectivity analyses. Resulting preprocessed data were utilized for subsequent analyses.

### 2.5 EEG source localization

To estimate the sources of the brain activity as reliably as possible, we implemented a source localization pipeline utilizing individual-level data. Each step is shortly described in the following paragraphs.

Head tissue segmentation of structural T1 images without the electrode net was performed by the automated 12-compartment segmentation tool (Taberna et al. 2021) which consists of image preprocessing, tissue probability mapping, and tissue segmentation steps. Subsequently, we followed a source localization pipeline based on the Fieldtrip toolbox (Oostenveld et al. 2011) functions. Based on all 12 compartments of the segmented T1, the hexahedral mesh was generated and compartment conductivities were assigned based on (Liu et al. 2018) to define a head model. A precise electrode localization for the head model, provided by Multimodal and Functional Imaging Laboratory of the Central European Institute of Technology (Brno, Czech Republic), was obtained semi-automatically from the acquired T1 image with the electrode net on, see subsection 2.2. Electrode locations were manually marked on rendered head surface where the electrode artifact was prominent. After the electrode location definition, structural T1 image with electrode net was coregistered to the second structural T1 without electrode net and resulting transformation matrix was also applied to the electrode positions. Electrodes were projected to the closest surface point of the head model. The source model was generated in the brain grey matter as well as cerebellar grey matter compartments with a grid size of 6 mm. The finite element method (FEM) SimBio (Vorwerk, Oostenveld, Piastra, Magyari, & Wolters 2018) solver was used to compute the leadfield matrix. The sources were estimated by the eLORETA inverse algorithm (Pascual-Marqui et al. 2011). Furthermore, we established a template source model utilizing the MNI-template anatomy from SPM (Penny et al., 2011) and the methodology stated above. This template source model was subsequently designated as the reference model for all subsequent analyses. Results obtained from the source space of the EEG-fMRI dataset, out-of-scanner EEG dataset, and fMRI were interpolated to this reference model using the nearest neighbor method. The source localization pipeline for out-of-scanner EEG data generally followed the same structure with the following differences: The source model resolution was 10 mm instead of 6 mm, the inverse warped source model grid from the MNI to the individual space was used instead of a regular grid directly generated in the individual space, the standard 5-layer head model in the Fieldtrip toolbox (Oostenveld et al. 2011) was used instead of the advanced 12-layer model in the MRTIM toolbox (Taberna et al. 2021), an electrode template was co-registered with an individual head structure and projected onto a head surface instead of individual electrode positioning, as discussed in section 2.1.2. Here, we followed a well-established Fieldtrip toolbox (Oostenveld et al. 2011) pipeline, mimicking a typical standard processing setup.

### 2.6 Electrode space spatio-spectral decomposition

In the present study, we assume that the electrical activity expressed as the EEG signal envelope measured at the electrode locations on the scalp is a linear mixture of source signals representing the brain activity. We further assume that the brain electrical activity power may have a different spatial profile across different EEG frequency bands, which is a substantial difference between the spatio-spectral models (allowing thus estimation of a much richer structure) and the trilinear models (Mareček et al. 2017; Marecek et al. 2016) where the spatial mode is assumed to be the same across the whole frequency range, only with a different strength. The schematic flowchart of the spatio-spectral decomposition methodology is depicted in Figure 1 . At first, we computed a signal envelope for each electrode and frequency band EEG time series by applying the Hilbert transform and then we downsampled the data to the fMRI BOLD TR (650 ms) (Brookes et al. 2011), obtaining thus a time series of band-limited power (BLP). Subsequently, we implemented an outlier correction procedure using Tukey’s fences (k = 3) on each time series. Outliers were identified and replaced with the mean value. Since the data were acquired in resting-state condition, we further band-pass filtered all BLP time series in a standard way to 0.008 - 0.09 Hz as for the resting-state fMRI BOLD data. Next, we concatenated the (space × time) BLP matrices of all frequency bands along the spatial domain, forming thus a joint spatio-spectral domain on the single-subject level. Each individual subject BLP time series was then z-score normalized to handle inter-individual variability. All single subject normalized BLP matrices were subsequently concatenated along the temporal domain forming a group BLP matrix *X*. Assuming that *X* is a linear mixture (by a mixing matrix *A*) of temporally independent BLP sources *S*, we can express the linear data generation process in a matrix form as:

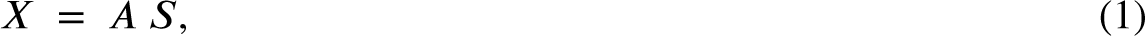

**FIGURE 1.**
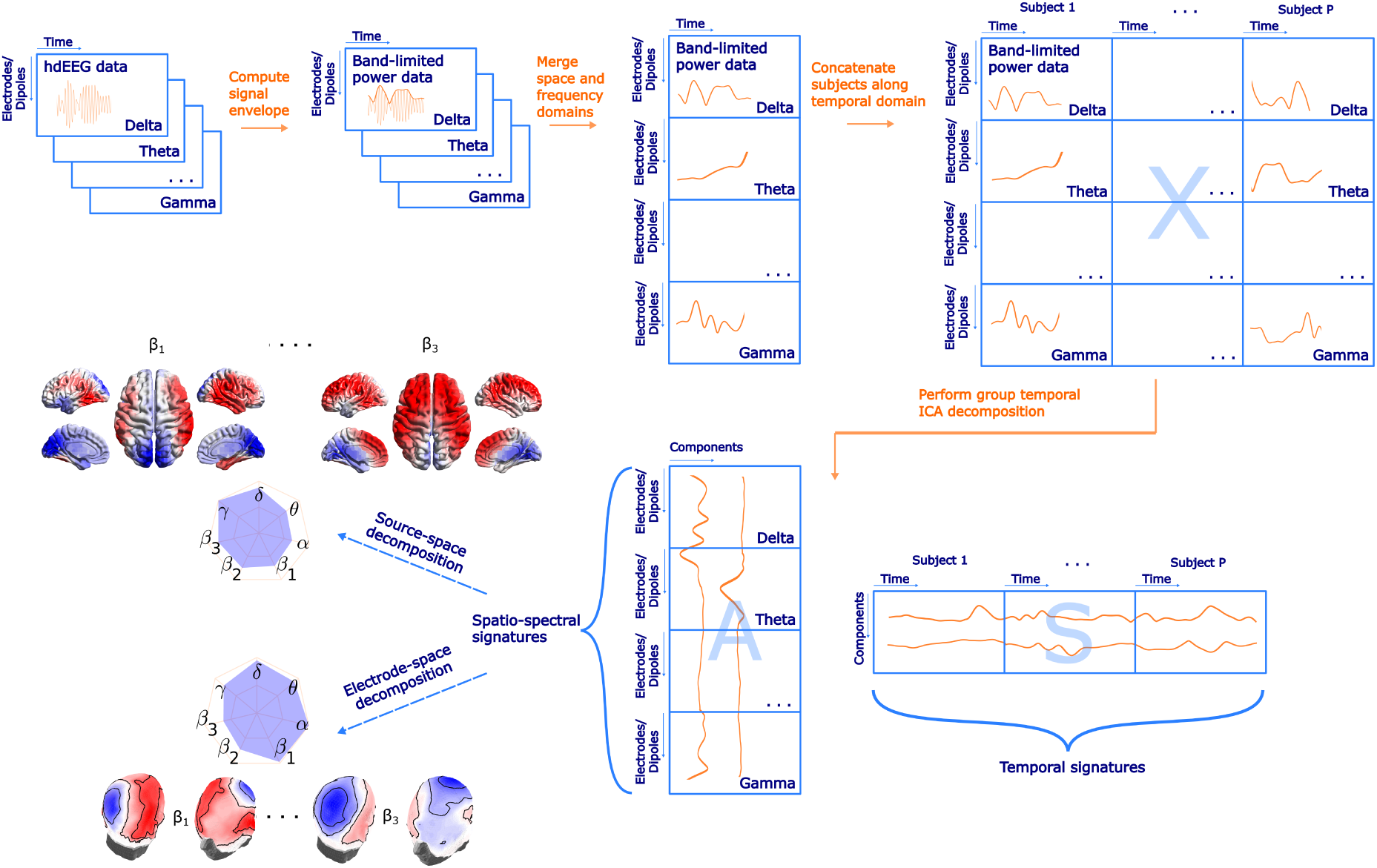
A schematic flowchart of the spatio-spectral decomposition in both electrode- and source space with highlighted main steps and examples of SSPs. Please note that electrode- and source-space data are subjected to the ICA decomposition separately. The label Electrodes/Dipoles is meant to denote in what dimension both approaches differ. Also note that the actual size of matrices in *Dipoles* dimension (*N_d_*, source-space version) is much higher compared to *Electrodes* dimension (*N_e_*, electrode-space version).

where the matrix *X* has size (*B* • *N_e_*) x (*P* • *T*). Here, *B* is the number of frequency bands, i.e., 7, *N_e_* is the number of electrodes, i.e., 195, *P* stands for the number of subjects, i.e., 72, and finally *T* is the number of single-subject BLP time points downsampled to TR, i.e., 1800. Applying a temporal (group-level) ICA by RUNICA algorithm (Makeig, Bell, Jung, & Sejnowski 1995) with a PCA dimensionality reduction set to *C* = 30 components. Several information criteria, including the Bayesian information criterion and the Akaike information criterion, were evaluated to ascertain the optimal number of components. Due to significant discrepancies between the criteria, the final number was chosen to align with the typical number of components in BOLD resting-state ICA analyses, see subsection 4 for further discussion. Thus, we obtain a matrix *S* with dimensions *C* x (*P* • *T*). Each row represents a single component time series considered to be a temporal signature of the given component. Matrix *A*, called *mixing matrix* with dimensions (*B* • *N_e_*) x *C* represents SSPs of the components (one in each column). Since ICA decomposition is based on iterative algorithms with random initialization, we utilized the ICASSO tool (Himberg, Hyvärinen, & Esposito 2004) to investigate the algorithmic reliability of the ICA decomposition itself by performing the ICA decomposition 20 times with different initial conditions. The cluster centroid time series were then considered as the most reliable estimate of components and therefore utilized for the subsequent analyses. Furthermore, SSPs were not obtained directly from the matrix *A* but via correlating each component time series (ICASSO cluster centroids) with each time series in BLP matrix *X*. The correlation coefficients were subsequently transformed by a Fisher Z-transformation. Obviously, each of the *C* = 30 SSPs consists of *B* spatial patterns (one for each frequency band), and each of *C* = 30 temporal signatures in *S* consists of concatenation of *P* individual time series (one for each subject). We also performed the spatio-spectral decomposition on the first and second half of the data set for stability analyses and statistical evaluation. To handle temporal discontinuities in the case of the out-of-scanner data set, we treated each data epoch separately up to a point before concatenation in the time domain. After that, the epochs within the subject were concatenated as well as subsequently across subjects. Finally, the ICA decomposition was performed in a similar way as for the previous data set.

### 2.7 Source space spatio-spectral decomposition

The source space spatio-spectral decomposition generally follows the workflow visualized in Figure 1 and described in sub-section 2.6 with the following differences specific to the source-projected EEG data. At each source model position, the dipole moment is expressed as three time series, one for each direction of the Cartesian coordinate system. Under the assumption that the direction of the dipole moment is not fixed but may rotate freely in time, we estimate the (scalar) signal power (amplitude) time series as in (Liu et al. 2017):

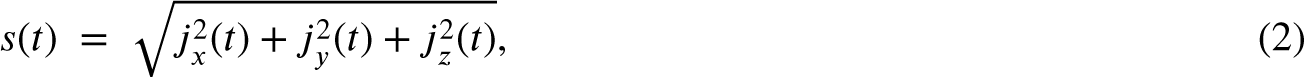

where *s*(*t*) is a BLP time series and *j_x_*, *j_y_*, *j_z_* are dipole moments in each of *x*, *y*, and *z* axes of the Cartesian coordinate system. Instead of *N_e_* spatial points or electrodes, in the source space case we obtain *N_d_* spatial points. These points correspond to the individual source model positions. As mentioned in 2.5, individual BLP time series were transformed to the template source model positions in order to allow concatenation across subjects. Decomposing the source BLP matrix enables us to spatially compare the SSPs with the statistical maps of the explained BOLD signal or with the BOLD-derived RSN spatial signatures. This was carried out by also interpolating the BOLD voxel positions to the template source model positions (rather than interpolating source model positions to the positions of the BOLD voxels), which was advantageous in terms of the computational complexity of the subsequent analyses. The out-of-scanner SSPs were also transformed into the template source model positions.

### 2.8 BOLD signatures of EEG spatio-spectral patterns

We reconstructed the EEG spatio-spectral component time series *S_p_* of individual subjects by simply splitting *S* to *P* segments of length *T* . To assess the single-subject relationship between EEG spatio-spectral component time courses and the BOLD signal in each voxel, the general linear model (GLM) was utilized. Note that several resting-state studies showed a substantial HRF variability of band-limited power regressor both when derived from a subset of electrodes (de Munck et al. 2007), using a bilinear (Labounek et al. 2019) or a trilinear (Marecek et al. 2016) band-limited power decomposition approach. To take into consideration such variability of the hemodynamic response function (HRF) across subjects, brain areas, and components, we included three regressors into the general linear model for each component separately: a component time series convolved with the canonical hemodynamic response function as the first regressor */J*_1_, and a component time series convolved with the first (temporal) and the second (dispersion) derivative of the canonical hemodynamic response function, as the regressors */J*_2_ and */J*_3_ – a procedure proposed and examined in previous event-related studies (Friston et al. 1998; Lindquist, Loh, Atlas, & Wager 2009). In contrast to (Labounek et al. 2019; Marecek et al. 2016), where authors employed F-statistics test inference for the GLM, our aim was to utilize such inference method specifically to discern the direction (positive/negative correlation) of the relationship between the component BLP time series and the BOLD signal. Therefore we implemented a formula proposed in (Calhoun, Stevens, Pearlson, & Kiehl 2004):

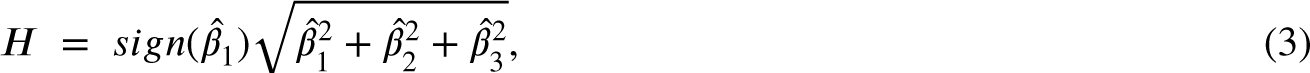

where *H* is an amplitude combining */J^r^*, */J^r^*, */J^r^* absolute values and the directionality is determined by the sign of */J^r^*, i.e., regression coefficient of component time series convolved with the canonical HRF. This combined voxel-wise beta coefficient *H* was subsequently used in a group-level analysis. Group-level statistics was performed by a one-sample t-test at each voxel and a cluster-based permutation statistics (Maris & Oostenveld 2007) was applied to correct for multiple comparisons (*α* = 0.05, *α_cluster_* = 0.05). Those fMRI BOLD activation maps (later referred to as BOLD signatures), as well as source space signatures, were visualized with the BrainNet Viewer tool (Xia, Wang, & He 2013).

### 2.9 Spatio-spectral EEG-fMRI integration evaluation

One of the challenges in the resting-state EEG-fMRI research is evaluating the reliability of the correspondence between both modalities. On one hand, there are methodological studies analyzing rather small EEG-fMRI data sets in terms of a number of subjects (Abreu et al. 2020; Sockeel et al. 2016; Yuan et al. 2016); thus, statistical power is low. On the other hand, there are studies examining reliability and reproducibility with a relatively larger amount of subjects (Bridwell et al. 2018) or across more data sets, and also across different paradigms (Labounek et al. 2019). Since we do not possess two equivalent EEG-fMRI data sets, we opted for checking the stability/reproducibility of observed patterns and relationships by splitting the sample - in our case we split our data set into two equal subsets, i.e., 36 subjects each. The choice of split into two equal subsets, was guided by practical considerations aimed at balancing statistical power with the need for subgroup analysis. This particular split allowed us to maintain a sufficiently large sample size in each subset, which is essential for observing typically weak levels of EEG-BOLD correlations while also facilitating comparisons between the two groups. We want to highlight that the methods described in the following parts can be broadly applied to other EEG-fMRI datasets acquired under different acquisition parameters. However, generalizing the findings and results presented in this paper should be approached with caution since they originate from one yet comprehensive EEG-fMRI dataset.

#### 2.9.1 Between-group stability of spatio-spectral EEG patterns

For an initial check of the robustness of the spatio-spectral patterns, we ran the SSP identification algorithm on each subset separately. We then computed Spearman correlation between each SSP from the first subset and each SSP obtained from the second subset, forming a square matrix with dimensions *C* x *C*.

A natural challenge in evaluation of the reproducibility of EEG or fMRI components is that the components provided have in principle arbitrary order, and are not thus directly comparable. This can be solved by sorting them by selecting the best match to each from a predefined list of templates (template-matching), or by simultaneously optimizing pairwise matches between all components of both subsets. We used this latter approach because it does not require selecting a template.

In particular, we first transformed this matrix by calculating absolute values and then multiplying by -1 to create a cost function for the Munkres algorithm (Bourgeois & Lassalle 1971), and then applied a MATLAB implementation of Munkres algorithm to find *C* unique pairs of SSPs between subsets. The algorithm satisfies a condition of maximum similarity and ensures that each component appears exactly once.

Note that establishing statistical significance for these maximum values is highly nontrivial, due to both the intricate sample dependencies, multiple testing, and ultimately the maximization procedure involved in selecting the best matches. Hence, we present the median and interquartile range of similarity in the results section. The results of the between-group stability of SSPs evaluation will be presented in subsection 3.1, along with the stability analysis of spatio-spectral EEG-fMRI integration. Indeed, while both the template matching and the global optimization procedures have previously been commonly applied to provide unique matching, the obtained best matches naturally suffer from upward bias related to the problem of overfitting by optimizing the matches, implicitly assuming a one-to-one mapping, that might be in many scenarios unrealistic, or carry biases in selection of the templates (if used). Thus, wherever practical, we developed alternative reliability testing procedures that avoid explicit template matching. In the following sections, we describe all statistical procedures used in this paper for evaluation of the proposed integration approach.

#### 2.9.2 Stability of spatio-spectral EEG-fMRI integration

While the reproducibility of the Spatio-spectral EEG decomposition *per se* is not straightforward to establish (see previous section), our main interest here lies in the EEG decomposition for the purposes of fusion with fMRI. Thus, in our evaluation, we focus rather on the stability of the link between an EEG component and its BOLD correlates, i.e. of the EEG-fMRI patterns. By EEG-fMRI pattern we mean the spatio-spectral and temporal signatures of a given EEG BLP IC together with the BOLD signature of this EEG BLP IC, i.e. the statistical GLM map obtained when using the EEG BLP IC time course as a regressor for concurrently measured voxelwise BOLD data. We work with the same subsets as in the previous case. For both subsets, a separate source as well as electrode space spatio-spectral decomposition was performed in the same manner as described in subsection 2.6 and 2.7, respectively. We test the hypothesis that components that have similar spatio-spectral signatures (between the test and retest decomposition), do also have a similar pattern in explained BOLD signal, i.e., BOLD signatures. To this end, a correlation matrix was computed between SSPs of the first and the second subset. Besides the correlation matrix of the whole length of SSPs, separate band-wise correlation matrices were also computed to test band-specific stability. After that, a group-level statistical GLM matrix having beta coefficients as columns for each component was created for both subsets, and a correlation matrix expressing similarity between components was computed. If the EEG components and their correspondence with the BOLD signal in both subsets are similar, then those two correlation matrices should be more similar than by chance, i.e. strength of match between the EEG SSPs should predict the strength of match between their BOLD signatures. To test this hypothesis, we implemented permutation statistical testing. During each of 1000 iterations, columns and rows of spatio-spectral correlation matrix were randomly permuted, and vectorized forms of the permuted SPP correlation matrix and of the GLM map correlation matrix were correlated to generate a permutation null distribution. Significance was determined based on a percentile of original not permuted similarity between spatio-spectral and statistical GLM correlation matrices (*α* = 0.05, right-sided test). Schematic flowchart of proposed testing of EEG-fMRI pattern stability is shown in Figure 2 . The results of this stability test are presented in subsection 3.1.

**FIGURE 2.**
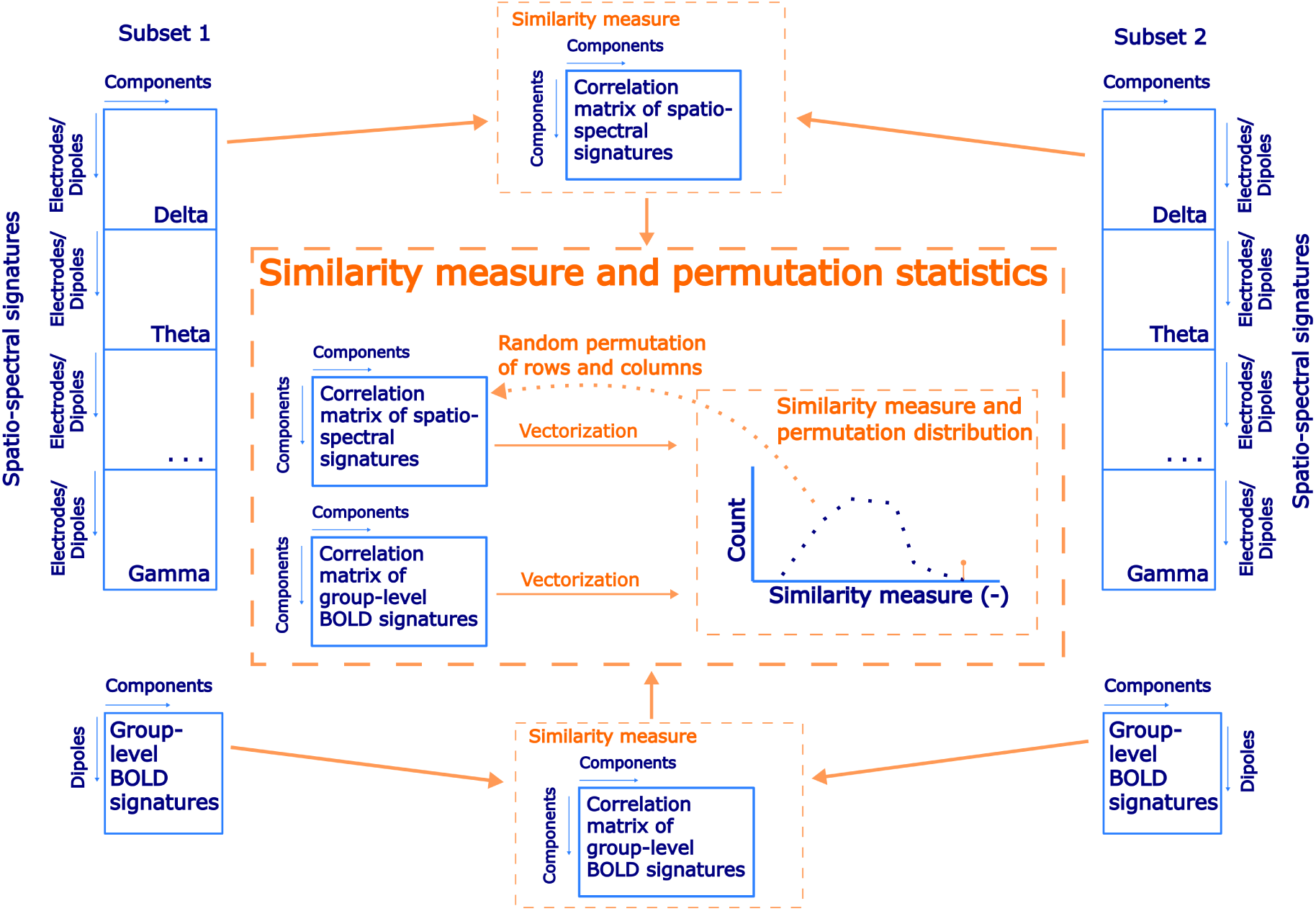
A schematic flowchart of a permutation statistical testing of EEG-fMRI patterns stability between subsets of the data set.

### 2.10 Identification of stable EEG-fMRI patterns

Following the assessment of overall stability in EEG-fMRI integration, our goal is to identify the most stable EEG-fMRI patterns across subsets that would call for more detailed neuroscientific interpretation. To achieve this, we introduce a heuristic metric designed to identify EEG-fMRI patterns that significantly contribute to the observed stability. This metric takes into account both the similarity of EEG BLP spatio-spectral maps and the similarity of the corresponding BOLD signatures. For each pair of spatio-spectral components within and across both subsets, we computed the following multiplicative criterion to quantify their similarity:

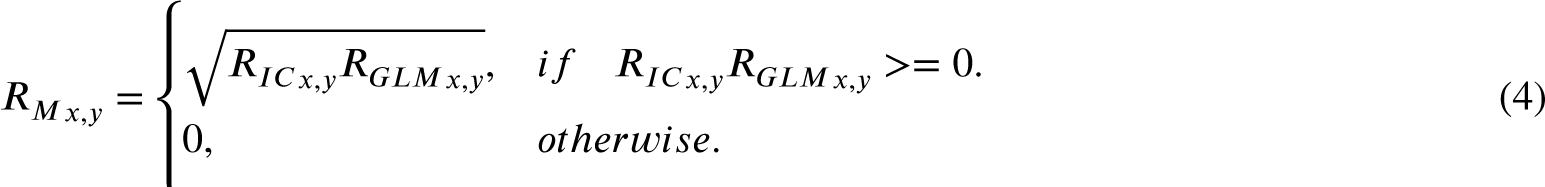

where *R_M x,y_* is the similarity (by multiplica^L^tive criterion) between components *x* and *y*, *R_IC x,y_* is the Spearman correlation coefficient between spatio-spectral signatures of components *x* and *y*, and *R_GLMx,y_* is the Spearman correlation coefficient between BOLD signatures of components *x* and *y*. This similarity metric was further transformed to a distance metric *D* = 1 − *R_M x,y_* for the application of a hierarchical clustering algorithm to form an agglomerative hierarchical cluster tree with the shortest distance method to compute a distance between clusters. We aimed to report only the first five most stable EEG-fMRI patterns (which corresponded to the threshold for defining clusters *D_thresh_* = 0.4 for both electrode and source spaces). To associate a single whole data set EEG-fMRI pattern to a pair of EEG-fMRI patterns from subset 1 and subset 2, we applied the same criterion as in equation 4 to both whole data set EEG-fMRI patterns with defined EEG-fMRI pattern from subset 1 and subset 2. The best-matching whole dataset pattern was determined as the one with the highest product value of these two criteria.

Up to this point, the stability testing and the identification of the most stable EEG-fMRI patterns was performed solely on the simultaneous EEG-fMRI data. To investigate whether the observed spatio-spectral components can also be found in the EEG data outside the MRI environment, we performed the following analysis. For each spatio-spectral pattern of the five most stable EEG-fMRI patterns, the most similar SSP in the out-of-scanner data set – based on the highest correlation of the spatio-spectral signatures – was determined and reported. It’s important to stress that reporting the most similar patterns from out-of-scanner data set does not provide a strict validation or generalization of the results to other data sets due to the discussed challenging problem of overfitting through the matching procedure. However, it demonstrates the level of variability to expect in resting-state EEG data.

### 2.11 Source space EEG patterns spatio-temporal relation to BOLD signatures and BOLD RSNs

So far, no one has directly spatio-temporally compared BLP-based features (in our case SSPs) obtained from EEG with their activation maps in BOLD (BOLD signatures). Even though it seems (at the first sight) natural that temporally related EEG and BOLD features should be also similar spatially, there is no strong evidence in the literature that this holds. Therefore, we propose two statistical testing procedures focused on: 1) purely spatial similarity of EEG SSPs and the corresponding BOLD signatures and 2) combined spatio-temporal similarity of EEG SSPs and the corresponding BOLD signatures.

#### 2.11.1 Spatial similarity of EEG spatio-spectral and BOLD signatures

Here, for the five most stable patterns we tested whether the similarity of their spatio-spectral signatures to their BOLD signatures is higher than to BOLD signatures of other EEG-fMRI patterns, indicating a potential spatial correspondence between spatio-spectral signatures and temporally coactivated areas observed in the BOLD data. To be able to compare the EEG spatio-spectral signatures and in this case purely spatial BOLD signatures, we separately correlated each band-wise part of SSP with a given BOLD signature and then computed a weighted average (weighted by the contribution of each band to the overall SSP weights) of the absolute values of Spearman correlations. In that manner, we computed a complete (weighted average) correlation matrix of all SSPs and BOLD signatures combinations. We compared the mean value on the diagonal (i.e. the correct correspondence between SSPs maps and BOLD signatures) to the null distribution obtained by computing the same statistic for each of 1000 random permutations of columns of the matrix, representing random assignment of different BOLD signatures to SSPs. The significance level *α* was set to 0.05 (right-sided test). We further evaluated for each component separately, how similar is to its corresponding GLM map compared to other BOLD signatures (each line of the correlation matrix). On the significance level *α* = 0.05 (right-sided test, uncorrected for a number of components), we report the most similar components. A schematic flowchart of the statistical testing is attached in the supplementary materials as Supplementary Figure S1. The results are presented in subsection 3.3.

#### 2.11.2 Spatio-temporal similarity of EEG spatio-spectral and BOLD signatures

Furthermore, we tested whether the EEG components more temporally synchronized with the BOLD signal tend to be also more spatially similar to their BOLD signatures. It is reasonable to assume, that multiple of the 30 EEG components may not correspond to strong stable neuronal dynamics, but rather to some transients or residual artifacts. To the extent that both BOLD and EEG reflect predominantly local neuronal activity (an assumption commonly used in brain activity modelling, based on experimental observations such as those presented by Logothetis et al. (2001) and others), one would expect that the stable EEG patterns would have BOLD signatures collocalized to the EEG spatial patterns, while the weaker, less robust, or artifactual components not temporally related to BOLD signal, would also have BOLD signature unrelated to the spatial EEG pattern.

To test this hypothesis, we use again a permutation-based test. In the same manner as in a previous subsection, we compute Spearman correlation between each of the 30 EEG spatio-spectral components and its BOLD signature, resulting in a vector. Then, for each EEG SSP, we correlate its time series with the BOLD signature time series. Since the BOLD signature is a statistical map representing spatial activation, it does not inherently possess a time series. Therefore, to establish a comparable time series for each BOLD signature, we computed the average BOLD time series. This average was computed using a weighted approach, where weights corresponded to the BOLD signature’s spatial pattern. By that, we prioritized voxels that showed a higher correlation with the EEG SSP time series in calculating this weighted average. This process also resulted in a vector of temporal similarities, with a length corresponding to the number of components, i.e., 30. If the described spatio-temporal similarity holds, these two vectors should be more similar than by chance. The null distribution was obtained by randomly permuting elements of both vectors (1000 iterations) and the original not permuted absolute value of Spearman correlation was tested against the permutation distribution at the significance level *α* = 0.05 (right-sided test). A schematic flowchart of the statistical testing is attached in the supplementary materials as Supplementary Figure S2. The results are presented along with the previous test results in subsection 3.3.

#### 2.11.3 Source space EEG patterns spatio-temporal relation to BOLD RSNs

In previous EEG-only studies (Liu et al. 2017 2018; Sockeel et al. 2016) and EEG-fMRI studies (Abreu et al. 2020; Marino et al. 2019; Yuan et al. 2016) the authors aimed to link EEG-derived RSNs and BOLD RSNs both spatially and temporally. We have decided to investigate spatial and temporal correspondence between EEG SSPs and BOLD RSNs involving a broad range of RSNs in the same unbiased spirit as for the previous tests. At first, we performed a spatial group ICA decomposition of the BOLD data. To this end, we concatenated all the single-subject z-score normalized datasets along the temporal domain. The dimensionality was then reduced to 30 PCA components before running spatial ICA decomposition by the FastICA algorithm (Hyvärinen & Oja 2000). Subsequently, ICs representing RSNs were identified based on functional network atlas obtained from (Shirer, Ryali, Rykhlevskaia, Menon, & Greicius 2012). The criterion was based on the Spearman correlation between the spatial distribution of the IC weights and atlas masks. Subsequently, statistical testing was carried out in an analogous fashion as for the spatio-temporal similarity of EEG SSPs and BOLD signatures. In particular, the (weighted across-band-averaged) correlation matrix of spatio-spectral signatures in the source-reconstructed space on one side with the spatial signatures of BOLD ICs spatial maps on the other side was computed. After that, temporal signatures of SSPs convolved with the canonical HRF were correlated with the BOLD ICs temporal signatures. If there was spatial and temporal correspondence between EEG-derived SSPs and BOLD IC components representing RSNs, then those two correlation matrices should be more similar than by chance. Similar to the approach in subsection 2.9.2, we implemented permutation statistical testing. In each of 1000 iterations, columns and rows of spatial patterns correlation matrix were randomly permuted (by two different permutations), and vectorized forms of the permuted spatial patterns correlation matrix and temporal patterns correlation matrix (absolute values) were correlated to generate a permutation null distribution. Significance was determined based on the percentile of the permutation distribution occupied by the original data similarity (Spearman correlation) between spatial and temporal patterns correlation matrices (*α* = 0.05, right-sided test). A schematic flowchart of statistical testing of EEG and BOLD ICs stability testing is in Figure 3 . The results are presented along with the two previous test results in subsection 3.3.

**FIGURE 3.**
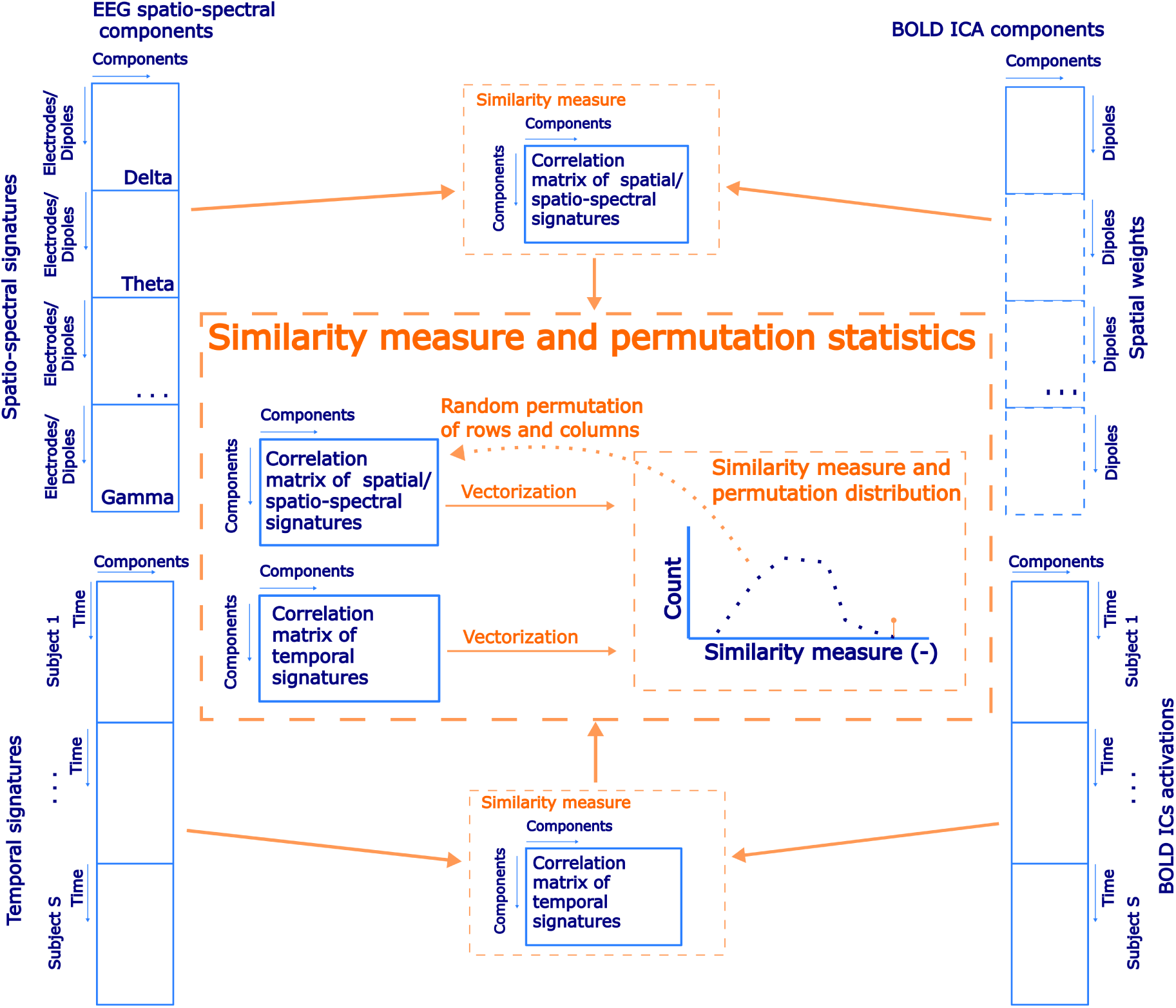
A schematic flowchart of permutation statistical testing of the correspondence of spatial and temporal similarities between the EEG source space spatio-spectral components and the BOLD ICs.

## 3 RESULTS

The results section is divided into three subsections. In subsection 3.1 we report EEG-fMRI patterns stability for source- and electrode space approaches, in subsection 3.2, we show the stable EEG-fMRI patterns in both spaces, in subsection 3.3, a summary from spatial and temporal correspondence testing of EEG SSPs, their BOLD signatures, and BOLD ICs is reported.

### 3.1 Stability of SSPs and EEG-fMRI integration

#### 3.1.1 Numerical SSP decomposition stability

As described in subsection 2.6, firstly, the spatio-spectral components stability testing in terms of ICA decomposition was performed utilizing the ICASSO tool. Based on a similarity index and visual assessment from the 2D CCA projection provided by the ICASSO, we conclude that the ICA decomposition is algorithmically stable for both source and electrode space approaches as well for both subsets and the whole data set. The ICASSO outputs are attached in the supplementary materials as Supplementary Figure S3.

#### 3.1.2 EEG-fMRI pattern reproducibility

The EEG-fMRI patterns proved generally reproducible in the test-retest setting, with the more matching EEG SSPs typically corresponding to those with more similar BOLD signature. This was true for both EEG spaces, with marginally better reproducibility for the source space approach, in particular the only two non-significant results were observed for the electrode space band-wise patterns in *8* (*r_s_* = 0.03) and *y* (*r_s_* = 0.04) bands. The highest reproducibility was observed for the multi-band (whole) SSPs - (*r_s_* = 0.29 for the source space and *r_s_* = 0.25 for the electrode space). The results are summarized in Table 1.

**TABLE 1.** Reproducibility of EEG-fMRI patterns: match of EEG SSPs corresponds to match of their BOLD GLM signatures. Table shows the similarity (Spearman correlation coefficient) of the correlation matrix of the spatio-spectral signatures obtained from the first and second data subsets with the correlation matrix of the respective BOLD signatures. Results are provided separately for the electrode- and source space approaches. ^*^ denotes permutation test significance (*a* = 0.05). All significant correlation coefficients remained significant also after FDR correction across tests (*a_FDR_* = 0.05).

| SSP | Spearman correlation coefficient (-) |  |  |  |  |  |  |  |
| --- | --- | --- | --- | --- | --- | --- | --- | --- |
| | whole | $\delta$ | $\theta$ | $\alpha$ | $\beta_1$ | $\beta_2$ | $\beta_3$ | $\gamma$ |
| Source space | 0.29* | 0.06* | 0.07* | 0.11* | 0.08* | 0.11* | 0.18* | 0.09* |
| Electrode space | 0.25* | 0.03 | 0.07* | 0.12* | 0.08* | 0.10* | 0.15* | 0.04 |

Of note, the SSPs stability test described in subsection 2.9.1 and based on components matching by Munkres algorithm clearly showed a *biased* result with median Spearman correlation 0.46 (IQR= 0.42, 0.58). This approach has two shortcomings: 1) It’s difficult to test the level of similarity between matched SSPs statistically, 2) It doesn’t test BOLD signatures stability. These results are further elaborated in the Discussion section 4.

### 3.2 Stable EEG-fMRI patterns

We identified the five most stable EEG-fMRI patterns based on the similarity measure and clustering described in subsection 2.9.2. The resulting hierarchical tree is visualized in Supplementary Figure S4. The stability is meant in the sense of similarity of SSPs and also of BOLD signatures. The five most stable source-as well as electrode space EEG-fMRI patterns are depicted in Figures 4 and 5, respectively (ordered based on descending similarity defined in subsection 2.10).

**FIGURE 4.**
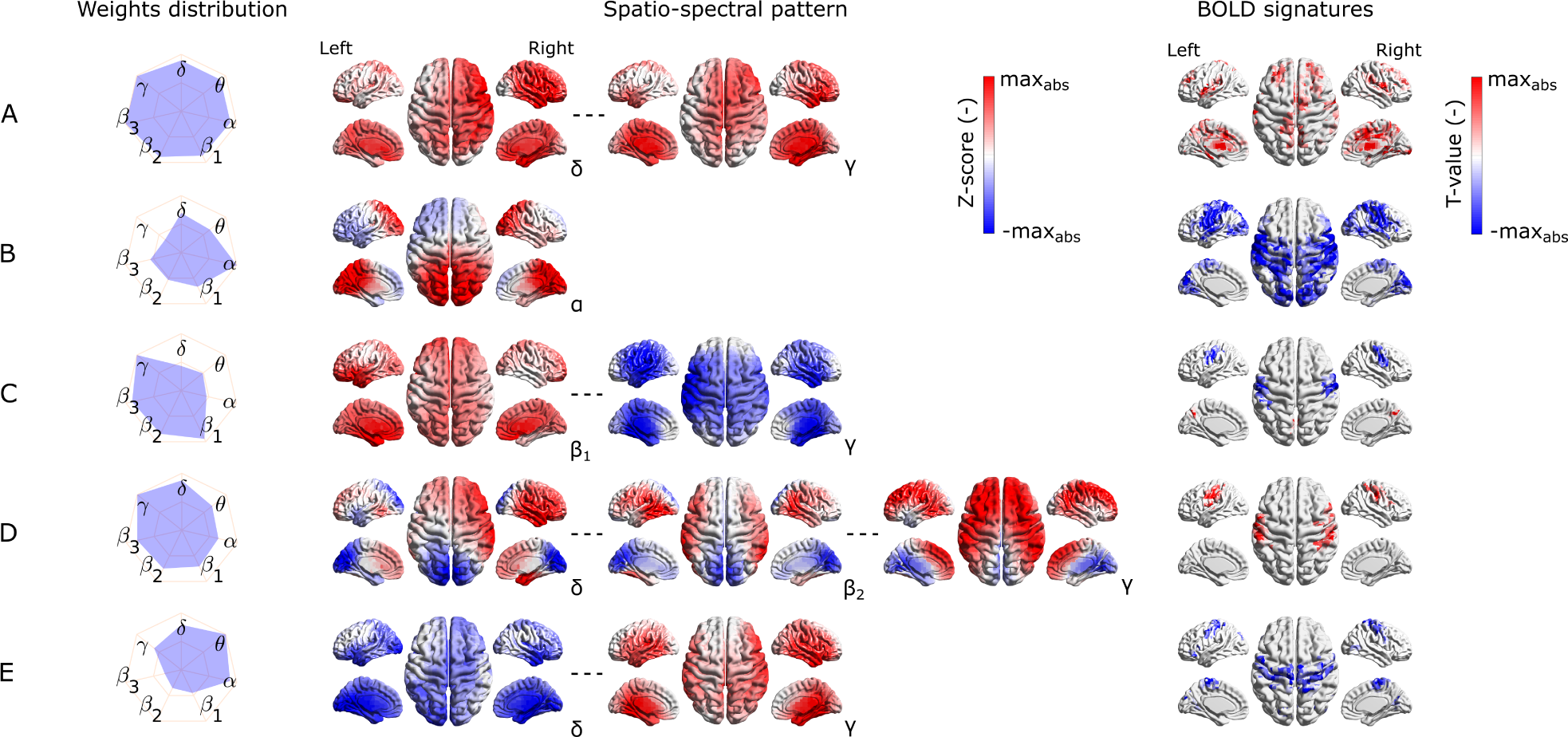
Visualization of the most stable source space EEG-fMRI patterns (A-E) in terms of the band-wise Euclidean norm of a SSP, SSPs, and the BOLD signatures (from left to right). For each EEG-fMRI pattern, only selected band-wise SSPs are visualized based on the weights distribution and their mutual similarity. Pattern **(A)** is similar between all frequency bands, pattern **(B)** is the most prominent in the *a* band, pattern **(C)** is similar in all */J* sub-bands and and has opposite weights in *y* band, pattern **(D)** in changes gradually from *8* to *y*) band. Finally, the pattern **(E)** changes its spatial distribution from *8* to *y* band.

**FIGURE 5.**
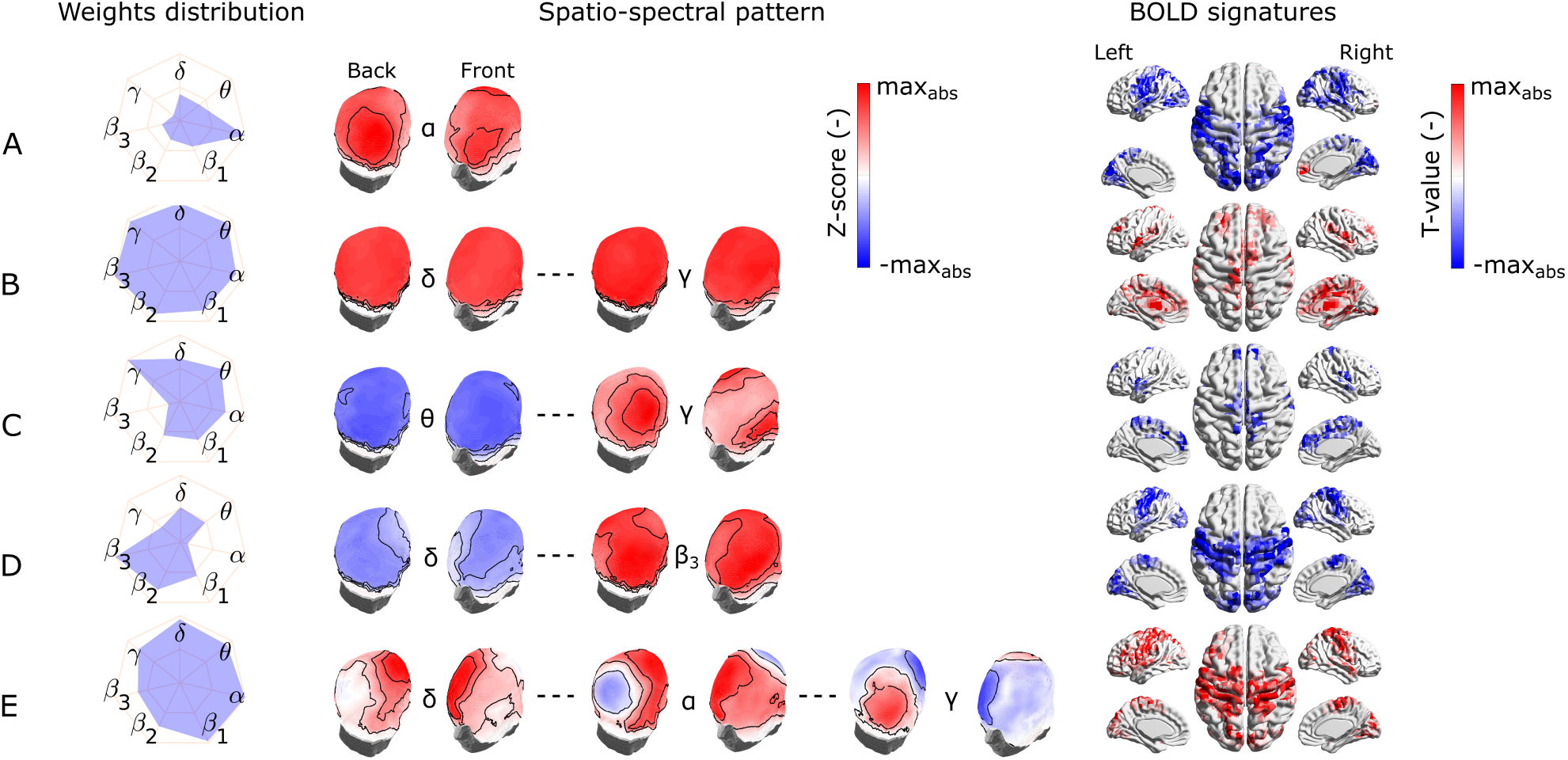
Visualization of the most stable electrode space EEG-fMRI patterns (A-E) in terms of the band-wise Euclidean norm of a SSP, SSPs, and the BOLD signatures (from left to right). For each EEG-fMRI pattern, only selected band-wise SSPs are visualized based on the weights distribution and their mutual similarity. Pattern **(A)** is the most prominent in the *a* band, pattern **(B)** is similar between all frequency bands, pattern **(C)** is similar for *0* band and neighbouring frequencies and differs in the *y* band, pattern **(D)** is less prominent in *8* and *0* bands and more in high (*/J*_2_ and */J*_3_) frequency bands. Finally, the pattern **(E)** changes its spatial distribution from *8* to *y* bands.

All these stable source as well as electrode space EEG-fMRI patterns have statistically significant correlation (cluster-based permutation test, *a* = 0.05, *a_cluster_* = 0.05) with the BOLD signal (see BOLD signatures in Figures 4 and 5). Moreover, they differ substantially in frequency as well as band-wise spatial distributions. Furthermore, some of the source space patterns are noticeably similar to electrode space patterns in terms of their BOLD signatures as well as ICs weights distribution.

#### 3.2.1 EEG-fMRI patterns common to source and electrode space

Specifically, the first EEG-fMRI pattern (Figure 4 A) is neither frequency-nor spatially specific and is represented by positive weights across all frequency bands and source positions, i.e. widespread positive weights from *8* to *y* bands focused more frontally and medially and less occipitally and parietally. This *global* pattern doesn’t correlate negatively with the BOLD signal but it shows positive correlation pattern with the BOLD signal in temporal (Heschl’s, superior temporal gyri), frontal (subcentral, frontal middle, frontal superior medial gyri, and supplementary motor area), parietal (supramarginal gyrus and precuneus), and occipital (lingual, occipital inferior gyri) lobes. Positive clusters also occupy insular and cingulate cortex and some subcortical brain areas such as amygdala, caudate nucleus, thalamus and putamen. In several parts of cerebellum (not visualized) positive clusters can also be observed. Noticeably, the second electrode space EEG-fMRI pattern (Figure 5 B) can also be called *global* since its weights distribution is broad and very similar across all frequency bands. This pattern correlates only positively with the BOLD signal in very similar brain areas as the mentioned pattern in Figure 4 A, although in contrast to the source space *global* pattern, the electrode space positive clusters were not observed in middle frontal gyrus and were in fusiform gyrus.

The following pattern (Figure 4 B) has weights distributed mostly around *alpha* band and spatially located positively across occipital, parietal and partly temporal lobes. This pattern has widespread negative correlation with the BOLD signal across the whole cortex and frontally focused positive activations. Negative clusters can be found almost over whole occipital lobe (calcarine, lingual, inferior, middle, superior gyri, and cuneus), parts of parietal (postcentral, supramarginal gyri, parietal superior cortex, and paracentral lobule) and temporal lobes (middle, superior temporal, Heschl’s, and fusiform gyri). In frontal lobe negative clusters are located in subcentral, precentral, inferior frontal gyri, supplementary motor area, rolandic and inferior frontal operculum. Positive clusters can be observed in anterior cingulate as well as parts of orbitofrontal cortices. This source space pattern is highly similar to the electrode space EEG-fMRI pattern shown inFigure 5 A. This pattern is spatially distributed over the whole scalp with the maximum in the parieto-occipital sites and also partly frontally. The statistical GLM map is again very similar to its source space counter part, only the electrode space component doesn’t have positive clusters in frontal lobe, while in some parts of cerebellum positive clusters were observed.

#### 3.2.2 Source space specific EEG-fMRI patterns

The third EEG-fMRI pattern (Figure 4 C) seems to be source space specific and is characterized by mostly fronto-parietal and medial positive weights across all three */J* sub bands and negative bilateral (mostly temporal) *γ* band negative weights. This pattern correlates positively as well as negatively with the BOLD signal. Negative cluster can be observed in parietal (postcentral, supramarginal gyri, inferior cortex, and paracentral lobule) lobe, temporal superior gyrus and in frontal (precentral, subcentral gyri, inferior operculum) lobe. Positive cluster is located in precuneus as well as again in several parts of cerebellum.

The next EEG-fMRI pattern (Figure 4 D) is characterized by negative parietal and occipital weights for all frequency bands and positive bilateral temporal and frontal weights from */J*_1_ to *y* bands. This pattern shows only negative BOLD correlate clusters in parietal (postcentral, supramarginal gyri, inferior cortex, and paracentral lobule) as well as in frontal (precentral, middle, superior, subcentral gyri, and inferior operculum) lobules.

Finally, the last EEG-fMRI pattern in the source space (Figure 4 E) is characterized by negative weights in low frequencies in bilateral temporal and occipital sites and at the same time positive *γ* weights again on bilateral temporal and occipital cortex sites. This pattern correlates exclusively negatively in parietal (postcentral, supramarginal gyri, inferior cortex, paracentral lobule and precuneus) and frontal (precentral, superior, middle and inferior gyri, supplementary motor area, and frontal inferior operculum) lobules as well as in middle part of cingulate cortex.

#### 3.2.3 Electrode space specific EEG-fMRI patterns

The third most stable electrode space EEG-fMRI pattern (Figure 5 C) is characterized by whole scalp negative weights in low frequencies around *0* band in contrast to positive weights in *γ* band spread mostly on parieto-occipito-temporal electrode sites. This SSP correlates positively with the BOLD signal in temporal (Heschl’s, superior and middle temporal gyri), frontal (supplementary motor area, roladic, frontal inferior operculum, frontal inferior triangularis, frontal superior, frontal superior medial and precentral gyri), parietal (precuneus, postcentral gyrus), and lingual gyrus in occipital lobe. Anterior and middle cingulate as well as insular cortices and parts of cerebellum also show negative correlation with the BOLD signal.

The fourth scalp SSP (Figure 5 D) is again not spatially specific and demonstrates changes in the band-limited power between low and high frequencies. Therefore, we observe gradually changing polarity of weights from the lowest frequencies *8* to */J*_3_ widely across the whole scalp. For this pattern specifically *a* and *y* frequency bands are the least important. The correlation pattern of the BOLD signal is exclusively negative and very similar to statistical GLM map of *α band pattern* (Figure 5 A). Apart from *α band pattern*, negative clusters can also be observed in parietal inferior, insular, middle cingulate cortices and frontal superior gyrus. A positive cluster is only in some parts of cerebellum.

The last electrode space EEG-fMRI pattern (Figure 5 E) is relatively complex. From *delta* to *theta* band the spatial pattern is mostly positive in frontal, central and temporal electrode sites. From *a* to */J*_3_ bands also negative parietal and occipital weights are present and *γ* band is represented by reversed pattern than for all other frequencies. This complex EEG-fMRI pattern has exclusively positive BOLD signature and it is very similar to previous negative BOLD signatures in Figures 5 A and D, respectively. In contrast to those, clusters in parietal and frontal lobes have relatively higher T values compared to the occipital ones.

#### 3.2.4 Variability of the five most stable EEG-fMRI patterns

Last but not least, to capture the variability of components between subsets of the same data set as well as between different data set recorded in a different environment (out-of-scanner data set), in Figures 6 and 7 we show the SSPs from both subsets and the closest SSP in out-of-scanner data set for each of five the most stable EEG-fMRI patterns for source-as well as electrode space approaches.

**FIGURE 6.**
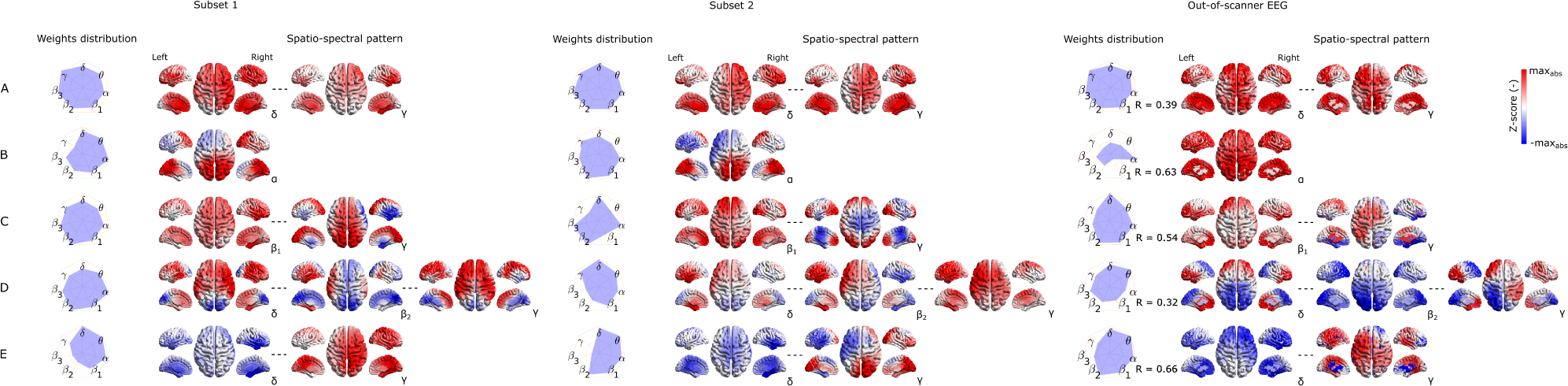
Visualization of corresponding subset 1 and 2 SSPs together with out-of-scanner SSP for each of the five most stable source space EEG-fMRI patterns (A-E).

**FIGURE 7.**
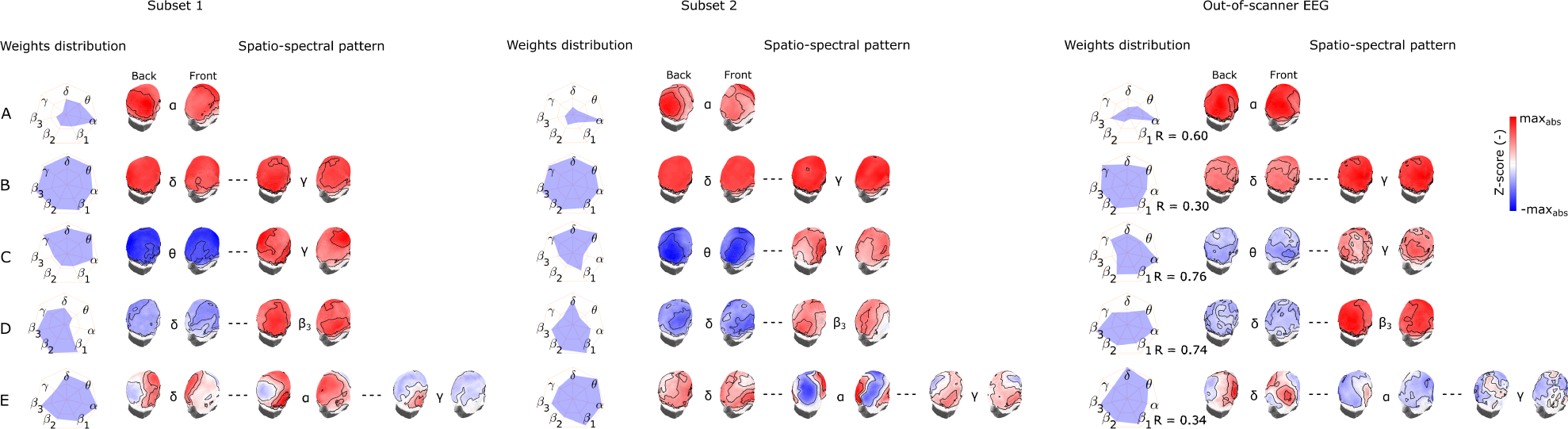
Visualization of corresponding subset 1 and 2 SSPs together with out-of-scanner SSP for each of the five most stable electrode space EEG-fMRI patterns (A-E).

Please note that in this case a statistical testing is not provided. Therefore both figures serve only as an example of SSP’s variability that can be expected when applying the methodology on a different data set. Also note that the most similar component of *a global* SSP from out-of-scanner data set was not determined based on the highest correlation with *a global* SSP from the whole data set but rather by selecting a component having all weights with the same sign. This feature can’t be measured with a similarity measure based on Spearman correlation. Also the visualized out-of-scanner SSP in Figure 6 C is the second most similar component to the whole data set component since the most similar component was the same as for Figure 6 E.

### 3.3 Source space EEG patterns spatio-temporal relation to BOLD signatures and BOLD RSNs

In this part, we study the spatial similarity of the EEG SSPs and the relevant BOLD structures, namely the corresponding BOLD signatures as well as independently constructed BOLD RSNs. The results are summarized in Figure 8.

**FIGURE 8.**
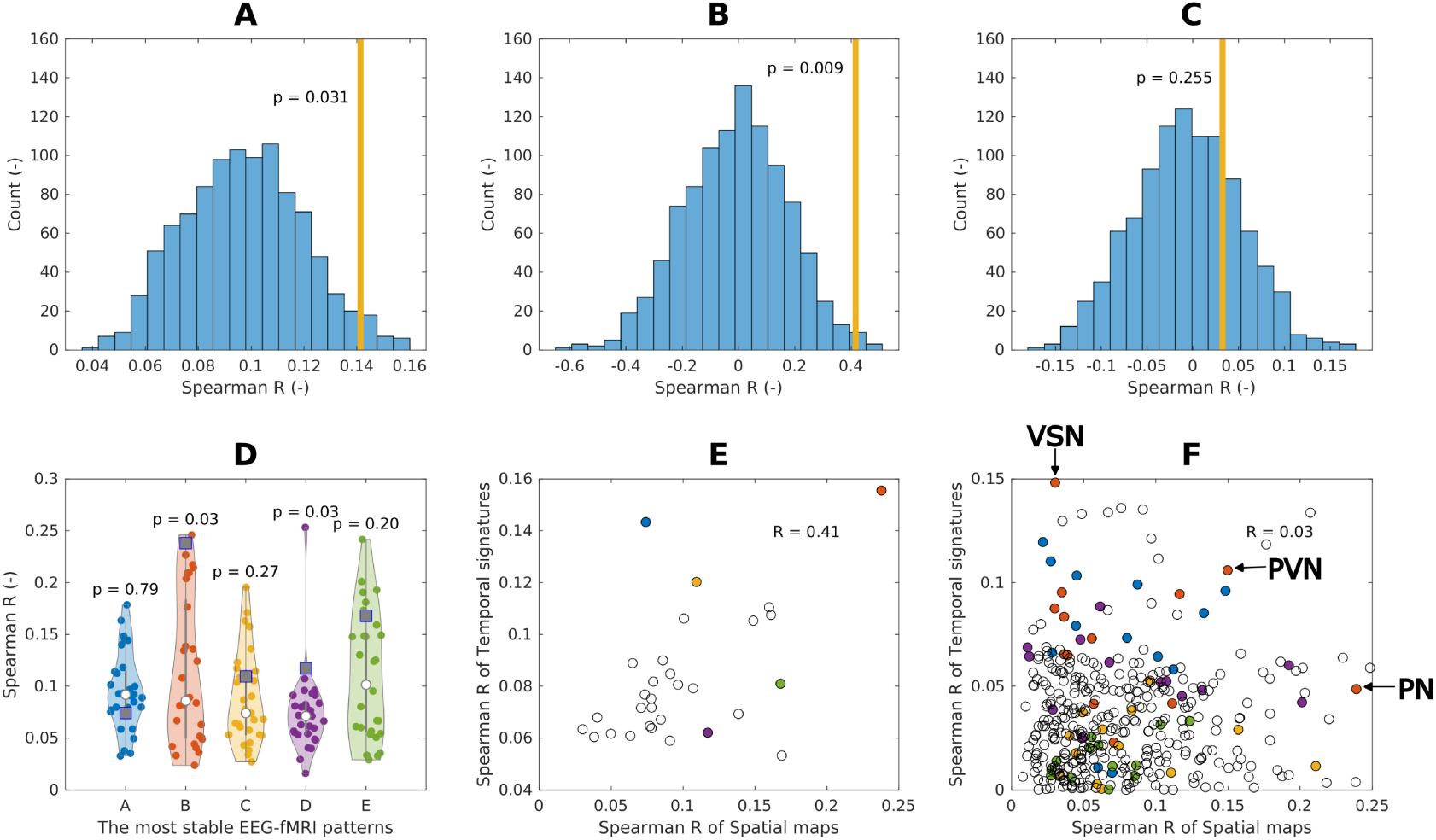
Spatial (spatio-temporal) relation of EEG spatio-spectral components and BOLD. **(A)** Spatial similarity of the EEG spatio-spectral signatures and their corresponding BOLD signatures: mean value across the five most robust components (orange bar), permutation test statistic (blue histogram). **(B)** Spatio-temporal similarity of the EEG spatio-spectral signatures and their corresponding BOLD signatures: mean values across all components (orange bar), permutation test statistic (blue histogram). **(C)** Spatio-temporal similarity of the EEG spatio-spectral signatures and BOLD RSNs: mean values across all components (orange bar), permutation test statistic (blue histogram). **(D)** Spatial similarity of all robust source space spatio-spectral patterns (A-E) all BOLD signatures (violin plots, dots) with denoted actual BOLD GLM map (square point). **(E)** Pair-wise similarity (Spearman correlation) of EEG spatio-spectral and BOLD signatures spatial (scatter plot x axis) and temporal (scatter plot y axis) signatures. Colored points correspond to EEG-fMRI patterns from **(D)**. **(F)** Pair-wise similarity (Spearman correlation) of EEG spatio-spectral and BOLD RSNs spatial (scatter plot x axis) and temporal (scatter plot y axis) signatures. Colored points correspond to EEG-fMRI patterns from **(D)**. Three points for *alpha* EEG-fMRI pattern are labeled as being: 1) most spatially similar to Precuneal network (PN), 2) most temporally similar to Visuospatial network (VSN), 1) most spatio-temporally similar to Primary visual network (PVN).

We found for the first 5 most stable EEG-fMRI patterns weak but significant (R = 0.14, p = 0.031) spatial relation between their spatio-spectral signatures and BOLD signatures, suggesting spatial colocalization between them, see Figure 8 A. In post-hoc testing, we observed that the similarity was mostly driven by a spatio-spectral pattern B and D, see Figure 8 B.

Suggested spatio-temporal test of EEG spatio-spectral components and BOLD signatures shows significant relation between the levels of their temporal and spatial correlation (R = 0.42, p = 0.009), suggesting that EEG spatio-spectral components having a higher level of correlation in the temporal domain with its statistical GLM map pair are also more similar spatially, see Figure 8 B and E.

Statistical testing of correspondence between EEG-derived spatio-spectral components and BOLD-derived ICs in terms of spatial and temporal similarities (R = 0.03, p = 0.255) didn’t reveal any similarity, see Figure 8 C and F. Specifically to demonstrate this on the most reliable *alpha* pattern, we picked 3 pairs of *alpha*-RSNs as: 1) The most spatially similar (PN), 2) The most temporally similar (VSN), 3) The most spatio-temporally similar (PVN) (up-right in scatter plot) pair *alpha* pattern-RSNs, see Figure 8 F. In Figure 9, the *alpha* EEG-fMRI pattern together with those 3 RSNs is visualized. These results are commented and put in the context in the Discussion subsection 4.

**FIGURE 9.**
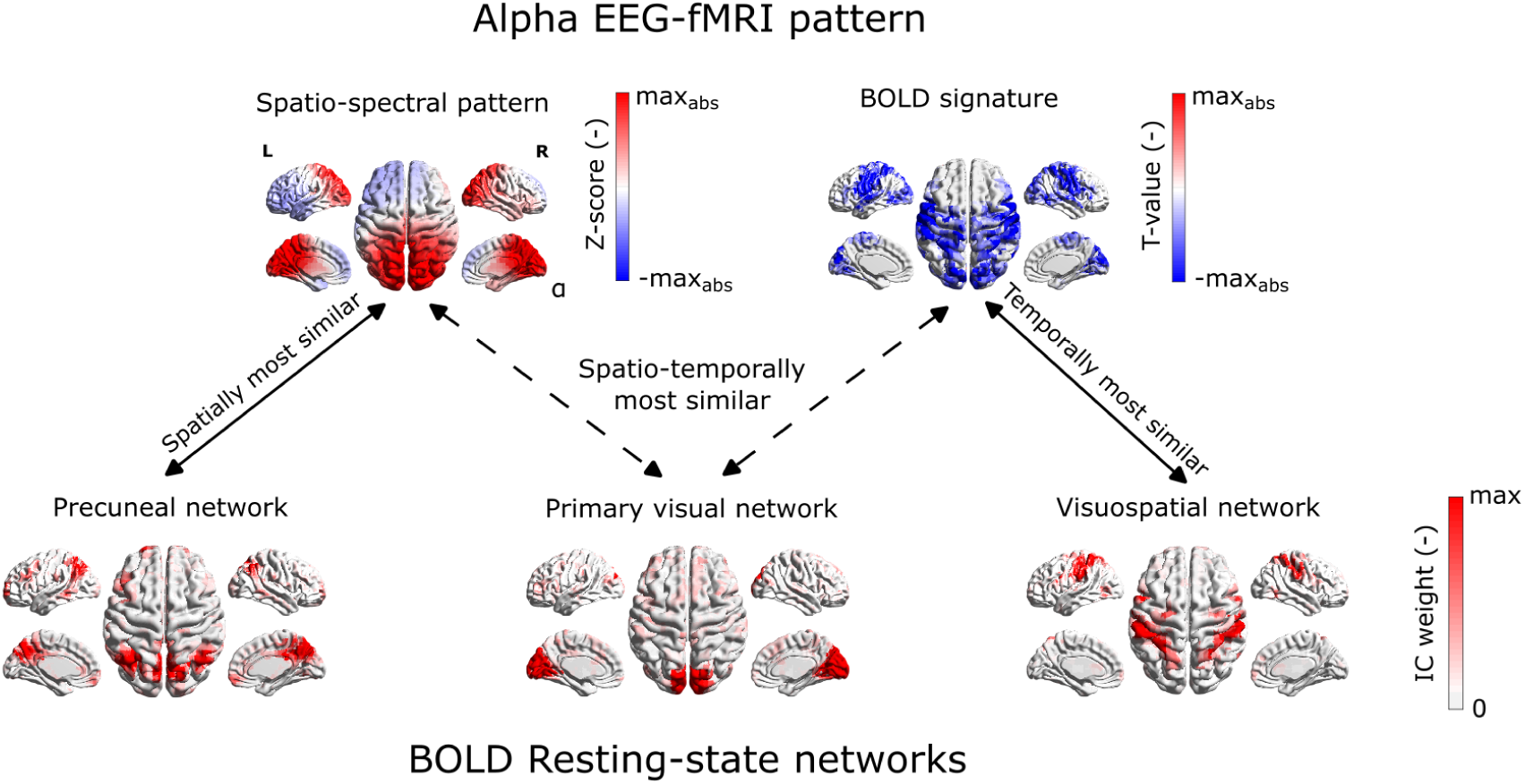
Example of spatial mismatch between alpha EEG-fMRI pattern and BOLD RSNs.

## 4 DISCUSSION

In the present study, we, for the first time, introduce a fully data-driven method of spatio-spectral decomposition in the source-reconstructed space applied to hdEEG data recorded simultaneously with the fMRI. We directly compare it to the electrode space spatio-spectral decomposition on the same data set. Rather than solely testing against other traditional methods, our study is focused on addressing intriguing questions regarding the EEG-fMRI relationship. Therefore, we apply a novel, unbiased approach to evaluate the stability of the observed EEG-fMRI patterns and, maybe even more importantly, of their spatio-temporal relation to their BOLD signatures and BOLD RSNs. A comparison with the SSPs obtained from the out-of-scanner data and with the results of electrode space decomposition is also provided.

### Stability of SSPs and their relation to fMRI

The unbiased stability testing based on the split-half analysis of the large EEG-fMRI data set clearly showed that the SSPs, and at the same time, their correspondence to the BOLD signal, are stable between subsets for both approaches (source and electrode space). All frequency specific (single-band) EEG spatial signatures are stable across subsets (besides electrode space *8* and *y* bands, see Table 1). The whole SSPs robustness suggests that the spatio-spectral components are formed across multiple frequency bands, as in the case of most electrode space (Figure 5 B-E) as well as source space (Figure 4 A,C-E) SSPs. Additionaly, (single-band) SSPs stability doesn’t exclude the possibility that some SSPs might be narrow (defined by a single frequency band) and that most frequency bands have at least a few unique components.

It is also interesting to note that the highest stability is achieved by the full (multi-band) spatio-spectral signatures compared to band-wise signatures for both spaces; see Table 1 . This suggests that the pattern across joint spatio-spectral space is important, and performing only single-band decomposition might miss some spectrally wide patterns.

In the presented work, the concurrent measurement with BOLD provided that we could simultaneously test SSPs and their BOLD signatures. Previously, when assessing the stability of EEG patterns between data sets or subjects, many studies utilized template matching procedures to pair the most similar components to BOLD-derived RSNs templates (Liu et al. 2017 2018; Marino et al. 2019) or component clustering algorithms applied only to EEG signatures (Labounek et al. 2019 2018). However, such matching approaches suffer from artificially inflated match quality (as optimal match is sought for across many components, even a random set of candidate maps, if sufficiently large, can provide decent ’optimal’ matches). For illustrative purposes and to maintain some comparability with results of previous studies (despite many other methodological differences), we also applied such straightforward matching procedure in the evaluation of SSPs reproducibility (the best matching SSPs between two subsets), where we reached a relatively higher mean level of similarity (*R* = 0.46).

A potential drawback of our stability analysis is the number of PCA components retained before the ICA decomposition, i.e., model selection. It’s because overestimating components may result in a component splitting (Abou-Elseoud et al. 2010). This could decrease the level of stability. It is difficult to adequately estimate the number of PCA components, and various model selection criteria often provide different estimates. Some authors estimate the number of PCA components on BLP matrices based on information theory criteria such as minimum description length (MDL) (Liu et al. 2017 2018; Marino et al. 2019). Others (Labounek et al. 2018) arbitrarily chose a number of ICs and tested the stability of recovered ICs by metrics obtained from toolboxes such as ICASSO software (Himberg et al. 2004) or setting the number of ICs to be around the typical number from fMRI studies (Li et al. 2018). A combination of the last two mentioned model selection strategies was utilized in the present study. A different approach might be to estimate the number of non-random components via a data-modeling and testing against surrogates (Vejmelka et al. 2015). Generally, the number of ICs throughout the literature fluctuates from 20 to 50; therefore, we chose the number of components in the current study to be 30. Another improvement might be a generalization of the stability evaluation between different data sets or even different paradigms, not only resting-state (Labounek et al. 2019 2018).

### Stable EEG-fMRI patterns

While we chose, in line with the literature, a generous number of 30 ICA components to be extracted, unlike several previous studies we present only a relatively modest number of 5 of them. Two factors may play a role here. Firstly, we pose an additional requirement on the presented components, that not only they have a counterpart in the decomposition of an independent EEG dataset, i.e. the EEG SSP patterns is reproducible, but moreover, we require that its BOLD signature is also conserved between the test and retest dataset. This focuses our analysis only on those EEG patterns that do have some robust BOLD signature, but perhaps even more importantly provides a control for false matches between random EEG patterns, which are likely to occur due to the sheer number of available components for the matching. Secondly, the fact that we look for multiband spectral patterns, that do not require the same spatial signature for each frequency, the hereby presented approach can hold richer patterns within one component, rather than splitting them into individual frequencies.

Thus, the resulting number of components described in detail in both source and electrode space is relatively smaller than those reported by other authors. Previous studies usually concentrated on single-band source space BLP decomposition and directly focused on spatial and temporal correspondence. They typically report almost a complete set of recovered EEG-derived RSN patterns. In EEG-only studies (Liu et al. 2017 2018), the authors reported finding a complete set of 14 EEG-BLP patterns spatially similar to BOLD-derived RSNs. Conversely, (Marino et al. 2019) focused on extracting a single DMN pattern from the source space EEG BLP matrix.

In (Bridwell et al. 2013), the authors reported ten robust EEG SSPs performing electrode space spatio-spectral decomposition. Five of those components were spectrally specific around the *α* band. Unlike our approach, the authors had a finer spectral resolution of 0.5 Hz, which, in combination with ICA decomposition along the spatio-spectral domain, may result in the decomposition of one alpha rhythm into several components. All five alpha components had notably similar occipital SSPs in their study, and they all correlated negatively with the BOLD signal in the occipital, parietal, and some parts of the frontal lobe. This would suggest that they may correspond to over splitting of a single relatively spatiotemporally consistent alpha component. The authors also report a widespread positive correlation across the cortex for the rest of the components, where three were in *8* - *0* and two in */J*_2_ - *y* frequency bands.

In (Labounek et al. 2019 2018), the authors found 14 SSPs (across three different data sets) having substantial similarity with components from (Bridwell et al. 2013). We report the five most reliable EEG-fMRI patterns, which are stable both in terms of SSPs and their BOLD signatures. Furthermore, it is noteworthy that some EEG-fMRI patterns between spaces (electrode and source level) intuitively correspond to each other. In the following paragraphs, we discuss all reported patterns in terms of interpretation and link to existing studies.

The most stable electrode space EEG-fMRI pattern (Figure 5 A) resembles the alpha components from a couple of previous studies (Bridwell et al. 2013; Labounek et al. 2018). It is a data-driven derived occipital alpha correlate known from many previous studies (Feige et al. 2005; Goldman et al. 2002; Gonçalves et al. 2006; Laufs, Kleinschmidt, et al. 2003; Laufs, Krakow, et al. 2003; Moosmann et al. 2003; Tyvaert et al. 2008). In agreement with (Bridwell et al. 2013), we found widespread negative correlation within a middle frontal, superior temporal, inferior occipital, lingual gyri, and cuneus, see statistical GLM map in Figure 5 A. Note that we also observed positive clusters in the anterior cingulate and parts of the orbitofrontal cortices. Positive correlations are not widely reported across studies except (Laufs, Krakow, et al. 2003), where the authors reported a positive correlation with the BOLD signal in the anterior cingulate cortex in a fixed effects group analysis. Intuitively, this electrode space component has its source space counterpart as EEG-fMRI pattern in Figure 4 B. This source space pattern has very similar statistical GLM map pattern corresponding to sensory areas and weights distribution, mostly in *α* band parieto-occipital cortical sites. Since the source space SSPs were not studied before, linking them with the current literature is difficult. Nevertheless, compared to the TFICA approach on EEG-only data set in (Li et al. 2018), this SSP may be related to the group of components called *Default* by the authors. It is also interesting that the BOLD signature of those components resembles a more typical EEG-fMRI occipital *α* band correlate rather than the bilateral frontoparietal pattern reported and discussed in (Laufs et al. 2006; Laufs, Krakow, et al. 2003).

The second pattern in the electrode space shown in Figure 5 B has (as in the previous case) its intuitive counterpart in the source space; see Figure 4 A. Those *global* patterns in both spaces also share very similar BOLD signatures when having exclusively positive clusters in all lobes and several subcortical structures such as thalamus, amygdala, or putamen. To further elucidate the existence of this pattern, it probably represents a global BLP mean time series since they appear to be correlated in space and across all frequency bands. This pattern may be notable in studies where authors derive BLP regressors as average BLP time series across all electrodes (Mantini et al. 2007). Our results show that the average BLP time series is one of the most stable (assessed by ICASSO) BLP independent components. Furthermore, we demonstrated that the average component belongs to a set of robust EEG components, significantly contributing to EEG-fMRI integration. Even though it is difficult to interpret such an unspecific EEG-fMRI pattern, this pattern might represent both neurophysiological activity and/or globally expressing artifactual patterns. The former (neurophysiological activity) explanation can be supported by the study (Magri, Schridde, Murayama, Panzeri, & Logothetis 2012). Despite substantial methodological differences (resting-state simultaneous LFP and BOLD data in monkey visual cortex), the authors made notable conclusions and remarks linking to this *global* as well as the previous *alpha* pattern. First, they found strong positive correlations between LFP at frequencies above 20 Hz and much weaker negative correlations between low-frequency LFP (less than 20 Hz) and the BOLD signal. Second, they report positive covariation of *α* LFP power and BOLD with total LFP power. Furthermore, they hypothesize that when significant fluctuations in total LFP power take place, the correlation between alpha power and the BOLD signal will seem positive (as reported by some authors), reflecting their shared variability with total LFP power. It seems that utilizing our method, the stable *alpha* pattern (anticorrelating with the BOLD signal) can be clearly distinguished from total power fluctuation (*global* pattern) allowing us to study those phenomena separately. Our method is therefore superior to ones considering only temporal averages of BLP across preselected electrodes/ROIs.

The next source space EEG-fMRI pattern in Figure 4 C represents an interplay between */J* and *y* frequency bands. The SSP for all */J* subbands can be described as primarily fronto-parietal medial positive weights compared to negative bilateral temporal and parietal pattern of the *γ* band. Notably, the BOLD signature of this pattern spatially resembles its SSP in terms of the positive cluster in the precuneus. Furthermore, parietal and temporal negative activations can be spatially linked to *γ* band as well. As the positive cluster in the precuneus (the central spatial component of the DMN) together with negative clusters (mainly SMN and AN functional networks), this SSP might represent temporal anticorrelation structure between DMN and so-called task-positive networks as suggested in (Fox et al. 2005) and later discussed in many other studies.

Another source space EEG-fMRI pattern in Figure 4 D reflects a broadband pattern from *8* to *y* band where negative weights can be observed through all bands in parietal and occipital lobes, whereas bilateral positive weights in temporal and frontal lobes are observed in higher */J*_1_ to *y* frequency bands. Based on its BOLD signature, this pattern might capture active somatosensory brain areas processing information; thus, positive correspondence between BOLD signal and */J*_1_ - *y* weights. At the same time, a decrease of BLP in parieto-occipital cortex sites might be associated with typical *α* band suppression and probably in combination with inactivity of visual processing since there is no positive cluster in occipital areas. Therefore, this pattern could be considered somatosensory specific and could distinguish between visual and other sensory processing.

The last not mentioned source space EEG-fMRI pattern relating low frequencies around *0* and *y* band is shown in Figure 4 E. Negative mostly occipital and midline low-frequency bands weights are included together with *γ* band bilateral frontal, temporal, and parietal positive weights. This pattern exhibits correlation with the BOLD signal in sensory areas such as postcentral and supramarginal gyri, parietal inferior cortex, and paracentral lobule. This pattern might, along with other EEG-fMRI patterns, reflect somatosensory processing.

The electrode space pattern in Figure 5 C also represents mutually anticorrelated BLP patterns between negative weights in low frequencies with a maximum around *0* band and positive *γ* band with a maximum in right parietal as well as left temporal electrode sites. Again, this pattern is not reported in previous studies. It is interesting to note that its BOLD signature differs from other EEG-fMRI patterns by having negative clusters in several parts of the frontal lobe, mainly in the superior frontal and medial superior frontal gyri, which might have a connection to typical sources of the midline theta rhythm (Asada, Fukuda, Tsunoda, Yamaguchi, & Tonoike 1999).

The EEG-fMRI pattern in Figure 5 D is again a frequency-wide but spatially not specific pattern showing a global anticorrelation pattern of BLP fluctuations between high and low frequencies. Interestingly, both *a* and *y* bands are the least important frequency bands for this pattern. This pattern is not observable in spatio-spectral decomposition studies (Bridwell et al. 2013; Labounek et al. 2019 2018). The rationale might be that it is unlikely that ICA decomposition performed along a spatio-spectral domain will retrieve such type of frequency-wide pattern. On the other hand, performing ICA decomposition along a temporal domain might reveal such types of spatio-spectral signature patterns. A more related frequency-wide component may be found in the trilinear PARAFAC decomposition, such as in (Mareček et al. 2017; Marecek et al. 2016), even though it is a different type of decomposition. Despite this component differing in spatio-spectral signature from SSP in Figure 5 A, the fMRI correlation pattern exhibits a significant negative relation when changing from high to low frequencies. This SSP also may be interpreted reversely such that large weights in lower frequencies *8* and *0* bands positively correlate with the BOLD signal across sensory brain areas in occipital and parietal lobes, which is in correspondence with low-frequency components in (Bridwell et al. 2013; Labounek et al. 2018).

Finally, the last not discussed electrode space EEG-fMRI pattern is a broadband and spatially specific pattern in Figure 5 E. Its SSP is represented by 1) frontal, central, and temporal positive weights for low frequencies; 2) *α* band negative parieto-occipital weights, and 3) reversed pattern in *γ* band. The statistical GLM map of this pattern is very similar (sign reversed) to one in Figure 5 A, except that clusters in parietal somatosensory areas have relatively higher T-values compared to occipital ones. Examples of SSP’s variability in Figures 6 and 7 show that to some extent similar components can be obtained not only from different subsets of the same data set but even from a different data set recorded out of the MRI scanner environment. Even though not statistically tested, this observation suggests that a similar methodology may also be applied to out-of-scanner data sets, which potentially increases the number of experiment types that the proposed methodology can investigate. Subjectively, *the global* and the *alpha* SSPs are the most robust for both source and electrode spaces. As highlighted in several sections of our manuscript, the out-of-scanner dataset differed from the simultaneous one in terms of how we acquired and analyzed the data. We acknowledge that these differences could have a meaningful impact on the obtained SSPs and, consequently, on the similarity levels of matched SSPs between the datasets. However, delving into a detailed exploration of the influence of these analysis differences is beyond the scope of our current study. This aspect might become a subject of investigation in future research.

In this work, we focus on the five most robust EEG-fMRI patterns. Indeed, further, particularly less prominent patterns could be extracted by alternative clustering setups (e.g., less strict thresholding) or a combination of dimensionality reduction and clustering algorithms such as applied in (Piorecky, Koudelka, Strobl, Brunovsky, & Krajca 2019). A particularly interesting directions would be to consider a possible hierarchical splitting of SSPs into more components and other aspects, such as linking between electrode space and source space components. Such an approach might also reveal differences in observed components between data spaces (electrode vs source) and that way help better understand a source space version of the spatio-spectral decomposition approach.

The use of source localization before the EEG power decomposition provides a direct anatomical link for spatio-spectral components, which we consider a major benefit of this method. In this study, a source space spatio-spectral decomposition approach did reveal source space unique patterns compared to the electrode space. We observed spatial correspondence between parts of SSP and its BOLD signature for some source space patterns. On the other hand, we should be aware of the ill-posed nature of the source localization and the resulting limitations of this technique (Hallez et al. 2007).

Last but not least, we want to make a comment on inter-individual variability. In this work, we applied a relatively conservative approach of the z-score normalization of each BLP time series. Nevertheless, there may be meaningful variability across subjects, as well as across bands/electrodes, that we here suppressed. This may be an objective of a future study.

### EEG spatio-spectral components relation to BOLD signatures and BOLD RSNs

Based on the results, we observed a far from perfect but significant spatial relation between EEG spatio-spectral components and BOLD signatures for the five most stable EEG-fMRI patterns. This finding is in line with the common assumption that the BLP fluctuations have spatially (almost) the same source as the BOLD signal. On the other hand, by looking at the most similar source space spatio-spectral signature and its BOLD signature, which is the *alpha* pattern (Figure 4 B), we can still notice that the spatial match is far from perfect. The similarity is mostly driven by both maps’ tendency to be more parieto-occipital rather than frontal. Besides all the potential sources of mismatch mentioned in the following paragraphs, the low levels of similarity can be caused by the ill-posed nature of the inverse problem. Indeed, one should consider that even in the case the fMRI and EEG power were perfectly collocalized, the EEG spatial distribution in the brain volume reconstructed from the electrode signals may achieve far from perfect recovery of this EEG-BOLD spatial match. The level of this reconstruction-based bias could be at least partially elucidated by simulation studies with realistic signals and forward and backward projection.

Furthermore, a significant linear relationship between temporal and spatial correspondence between EEG spatio-spectral patterns and their BOLD signatures further suggests that there exist EEG components more informative for the BOLD signal, which is (at the same time) more spatially similar to their BOLD signatures. Suppose we assume that this correspondence arises from physiological brain activity. In that case, it means that the more of the same *brain activity* visible in both EEG and BOLD, the more spatially similar they are and vice versa. On the other side, our statistical evaluation could not confirm the spatial and temporal correspondence between BLP-derived components and the BOLD ICs. This negative finding might contrast with studies (Liu et al. 2017 2018; Marino et al. 2019). Via a group ICA approach, obtained components represent the structure of the whole data set. Of note, our approach does not require any pattern matching of components between subjects, and avoids thus potential overfitting.

Our approach is of course vulnerable to the ubiquitous problem of selecting the number of ICs recovered and the previously mentioned problem of component splitting, which may cause a problem for the selected similarity statistical method. Nevertheless, our study suggests that while one may find for known BOLD ICA components some EEG SSP patterns that would be spatially similar (as shown previously in the literature), there is not a clear spatio-*temporal* agreement between *main components* of EEG BLP and the BOLD ICA components. More research is thus needed to clarify the spatial (spatio-spectral) and temporal correspondence between EEG BLP patterns and BOLD ICs.

For an illustrative example, as demonstrated in Figure 9, we see that even for the most stable EEG-fMRI *alpha* pattern (based on several previously described criteria), there is no clear RSN counterpart that would be the most similar spatially and at the same time temporally. Furthermore, those related networks (Precuneal network, Visuospatial network, Primary visual network) are associated with a different type of brain processing. Therefore, when performing an EEG-only study, we should be careful when interpreting the resting-state results based on only spatial profiles of BLP features in the sense of BOLD functional networks. In any case, the direct spatio-temporal relation of EEG BLP and RSNs does not seem trivial, and one should keep in mind that we cannot use knowledge from one modality to interpret results from other modality solely on *spatial colocalization* assumption and without further discussion.

The statistical methods used to test the stability and reliability of EEG-fMRI patterns, as well as the relationship between EEG SSPs and BOLD signatures/RSNs, involved multiple levels of construction and processing of descriptive statistics that were only at the end subjected to statistical hypothesis testing. This resulted in an increased abstraction level of the final results. For instance, when testing the *spatial colocalization hypothesis* in subsection 2.11.1, we began with derived EEG spatio-spectral patterns, which are ICs representing band-limited power fluctuations in EEG. We believed that the spatio-spectral signature of a given component represents a spatial pattern (one for each frequency band), and its temporal signature depicts the evolution of this spatial pattern over time. To obtain a BOLD-related map, it’s straightforward to use a temporal signature of a given EEG component as an explanatory variable in a typical GLM analysis as is subsection 2.8 and obtain BOLD signatures (a spatial map of beta coefficients). In the ideal case of *spatial colocalization*, the BOLD signature as a spatial map would perfectly spatially correspond to the spatio-spectral signature of a given EEG component. We can compute the level of spatial similarity of those two maps by (in our case) Spearman correlation. To address the issue of separate spatial maps for each EEG frequency band, we applied a weighted average. Additionally, given that both types of spatial maps (EEG SSPs and BOLD signatures) exhibit spatial smoothness, assessing the level of statistical significance was challenging. Therefore, as a final step, we employed permutation statistics to generate a meaningful null distribution. In summary, by the *spatial colocalization*, we tested whether the spatial similarity between a given EEG SSP and its BOLD signature (assessed by the weighted average of band-wise spatial map similarity with BOLD signature) was, on average, significantly higher than between the same EEG SSP and BOLD signatures of other components. In other cases, as in subsections 2.9.2, 2.11.2, and 2.11.3, we utilized a very similar methodology, customizing this multi-step approach to suit each specific hypothesis. Our aim was to consistently compare comparable derived measures (spatial against spatial and temporal against temporal) and to construct a meaningful null distribution for statistical analysis. Of note, the thresholding of the various spatial maps prior to statistical analyses was omitted because those analyses focused on comparing overall pattern similarity between the maps rather than interpreting individual map values. This omission did not introduce bias into the analysis but might introduce some level of noise.

A possible explanation for not observing spatial/spatio-spectral and temporal correspondence between EEG BLP patterns and BOLD ICs might lie in brain activity phenomena that are assumed to give rise to these two signals. While EEG signal is considered to rise from temporally and spatially synchronous neuron activity, specifically excitatory post-synaptic potentials/currents, the BOLD signal generation is dependent on the rather complex vasodilation effect, which is controlled by multiple different pathways as summarized in (Drew 2019). It is also a matter of ongoing research which (and how much) cell types in the central nervous system (CNS) contribute to the overall captured BOLD signal. In an optogenetic mouse model stimulation study (Vazquez, Fukuda, & Kim 2018), the authors found that stimulating interneurons leads to a larger increase in blood flow than stimulating pyramidal neurons; thus, they hypothesize that interneurons might be the primary driver of the BOLD response. Other mouse model studies suggest that also astrocyte responses might contribute to the BOLD signal (Tran, Peringod, & Gordon 2018) and that astrocyte activity might not modulate neuronal activity (Takata et al. 2018). In contrast to human resting-state EEG-fMRI studies, in many animal studies, it was observed that typically gamma band-limited power (Logothetis et al. 2001; Mateo, Knutsen, Tsai, Shih, & Kleinfeld 2017; Schölvinck, Maier, Ye, Duyn, & Leopold 2010; Winder, Echagarruga, Zhang, & Drew 2017) and multi-unit spiking activity (Ma et al. 2016; Mateo et al. 2017; Winder et al. 2017) correlates with the BOLD signal. It is also discussed in (Drew 2019), that observed low correspondence between EEG and fMRI resting-state signals (correlation coefficient typically lower than 0.3) might not be caused by methodological issues but rather by ongoing vasculature dynamics that might be independent of neural activity. This is argued by evoked activity studies (Lima, Cardoso, Sirotin, & Das 2014; Logothetis et al. 2001; Winder et al. 2017), which report much higher levels of correlation (even higher than 0.9), thus a high level of explained variance. Findings in those animal studies, as well as other non-neural signals that are superposed on *true* neural activity and hemodynamic responses, might all contribute to not directly observing spatio-temporal relation between the proposed EEG- and BOLD-derived patterns. These findings and theories follow mostly the assumption that EEG- and BOLD-derived fluctuations are spatially colocated. Nevertheless, since generators of EEG are generally considered EPSCs, and we assume long-range communication between brain areas, the active brain area (observed in BOLD signal) causes observable EEG activity at a different part of the brain. Therefore perfect spatial colocalization of EEG and fMRI might not likely emerge. In that case, studying causal relationships might further contribute to a better understanding of those modalities. It’s also important to note that there are other EEG features distinct from the amplitude of band-limited EEG oscillations relating EEG and BOLD signals such as reported BOLD correlates of infraslow EEG fluctuations (ISF) Hiltunen et al. (2014); a signal that is known to be correlated to amplitude of band-limited EEG oscillations Monto, Palva, Voipio, and Palva (2008), although the relation of the BOLD correlates of ISF EEG and BLP power oscillations has not yet been explicitly studied. Finally, we must also acknowledge the impact of EEG data quality acquired within the MRI environment. Despite employing state-of-the-art methods for EEG preprocessing, effectively suppressing the two most prominent artifacts (pulse and gradient) remains challenging, partly also due to their mutual interaction (Steyrl & Müller-Putz 2019). One such improvement of the proposed methodology would be to evaluate the quality of EEG recorded during fMRI with the in-scanner EEG recorded prior to fMRI acquisition. Further development of the preprocessing methods is necessary primarily because the mentioned artifacts partially overlap with the EEG frequency bands of interest.

## 5 CONCLUSION

We comprehensively compared and integrated EEG whole-brain patterns of neural dynamics with concurrently measured fMRI BOLD data. For that purpose, we derived EEG patterns based on a spatio-spectral decomposition of band-limited EEG power in the source-reconstructed space. The decomposition has proven stable in terms of the similarity of the EEG spatio-spectral signatures with the correlation patterns of the BOLD signal, and we further illustrated that the SSPs obtained from the source-reconstructed space are anatomically interpretable. Furthermore, we observed statistically significant but weak spatial correspondence between EEG spatial profiles and their BOLD signatures, supporting the plausibility of both in terms of mutual validation, while challenging the view of perfect spatial co-localization of EEG power and BOLD fluctuations. Moreover, we did not observe substantial spatiotemporal correspondence between the EEG and BOLD components. Overall, the introduced data-driven method with enhanced spatial resolution may be useful for a deeper understanding of the link between EEG and fMRI, namely human brain networks. Specifically, this approach might identify complex spatiospectral dynamics of interest not only in rest, but also tasks studies of the dynamics of brain activity both in health and in disease states.

From the data analysis perspective, we showed that EEG spatio-spectral decomposition in the source-reconstructed space is a viable dimensionality reduction technique that provides direct information about an anatomical region related to a specific spatio-spectral pattern and, therefore, provides EEG components that are directly spatially comparable to BOLD spatio-temporal patterns. The presented results shed more light on resting-state EEG-fMRI literature when understanding the link between those two modalities. From an application perspective, the extracted EEG spatio-spectral patterns may serve as a spatio-spectral filter for various applied studies (i.e. provide well-informed dimension reduction to time series of 5 interpretable SSP patterns) or as a basis for other methodological EEG-fMRI work studying the link between those modalities.

To further explore the spatio-spectral space by ICA or similar methods, advanced clustering analysis may contribute to potentially detecting subsequent – weaker or more detailed – patterns. Different types of more subject-individual decomposition methods might be beneficial in understanding more specific patterns. A good trade-off between single-subject ICA decomposition and the group ICA method presented in this direction might be an independent vector analysis (IVA) (Lee, Lee, Jolesz, & Yoo 2008). There also exist other methods from the trilinear decomposition family, such as Tucker trilinear decomposition methods (Tucker 1966) or block-term decomposition (Rontogiannis, Kofidis, & Giampouras 2021). Those are far less constrained approaches (compared to PARAFAC) which might provide valuable insights into interactions within EEG data. The proposed methodology may also serve for computational modeling purposes as an EEG feature extraction method for whole-brain network computational models (Cakan, Jajcay, & Obermayer 2021; Sanz Leon et al. 2013; Schirner, McIntosh, Jirsa, Deco, & Ritter 2018).

## AUTHOR CONTRIBUTIONS

SJ set the overall structure of the paper as well as analyses structure and he also provided analyses on both EEG and fMRI data. VK provided his expertise in EEG source localization analysis and set the overall structure of the paper and supported data visualization. DM provided his expertise in EEG-fMRI integration and provided the software for EEG preprocessing. RM provided his expertise in EEG-fMRI integration, blind source separation methods, and provided the data set and also he preprocessed the fMRI data. JH secured resources and supervised the whole study, with particular contribution to the design of EEG-fMRI integration methods and statistical analyses design.

## ACKNOWLEDGMENTS

The publication was supported by ERDF-Project Brain dynamics, No. CZ.02.01.01/00/22_008/0004643, the Czech Technical University Internal Grant Agency - grant number SGS23/120/OHK3/2T/13, Ministry of Health Czech Republic - DRO 2021 (”National Institute of Mental Health - NIMH, IN: 00023752”), and the Czech Science Foundation project No. 21-32608S. We acknowledge the core facility MAFIL supported by the Czech-BioImaging large RI project (LM2018129 funded by MEYS CR) for their support with obtaining scientific data presented in this paper. We would like to express our gratitude to Marco Marino, Gaia Taberna, and Jessica Samogin for their contributions to EEG preprocessing and forward modeling software development.

## DATA AVAILABILITY STATEMENT

The data that support the findings of this study are available from the corresponding author upon reasonable request.

## SUPPLEMENTARY MATERIALS

Supplementary figures (S1 - S4) are provided within a separate supplementary file.

