## Supplementary figures for "Spatial (Mis)match Between EEG and fMRI Signal Patterns Revealed by Spatio-Spectral Source-Space EEG Decomposition"

### Supplementary material

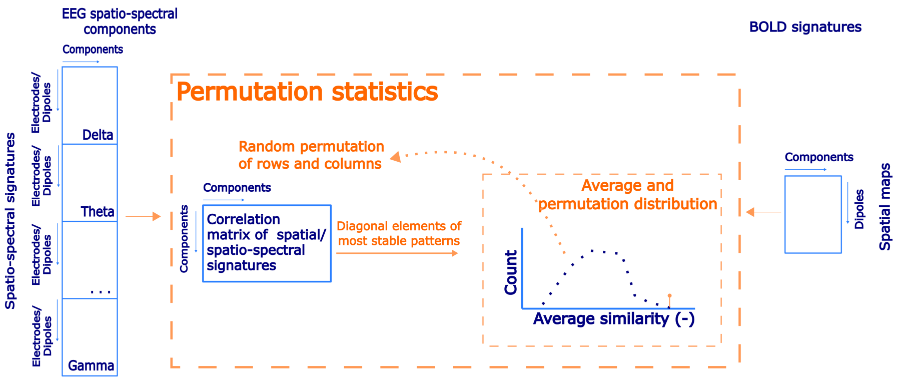


Figure S1: A schematic flowchart of a permutation spatial statistical testing of EEG spatio-spectral signatures and statistical GLM maps.

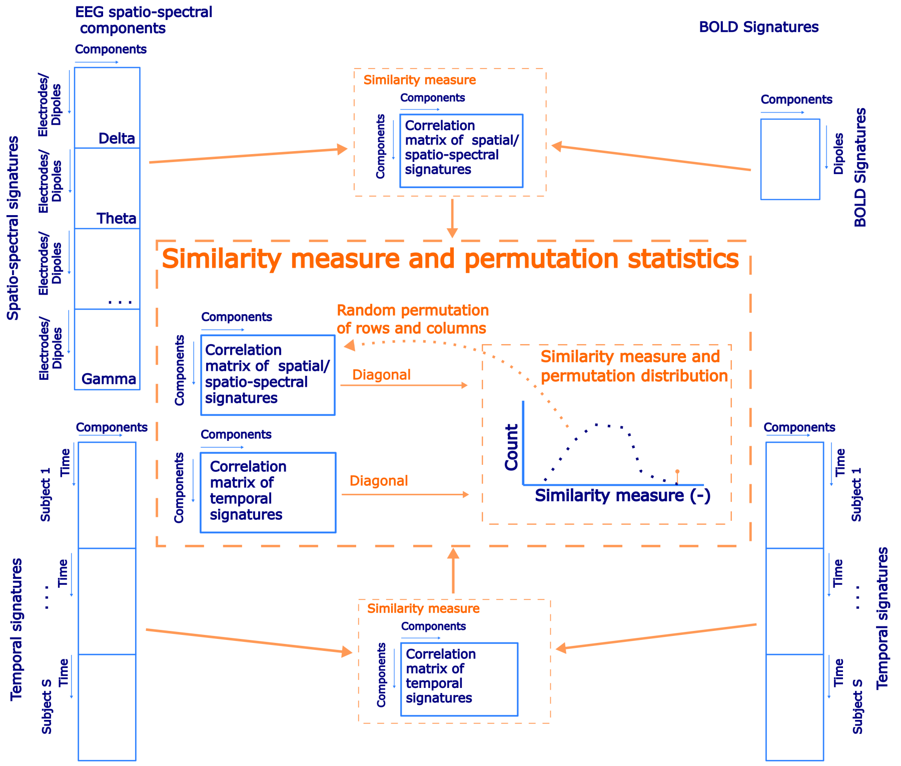


Figure S2: A schematic flowchart of a permutation spatio-spectral statistical testing of EEG spatio-spectral signatures and statistical GLM maps.

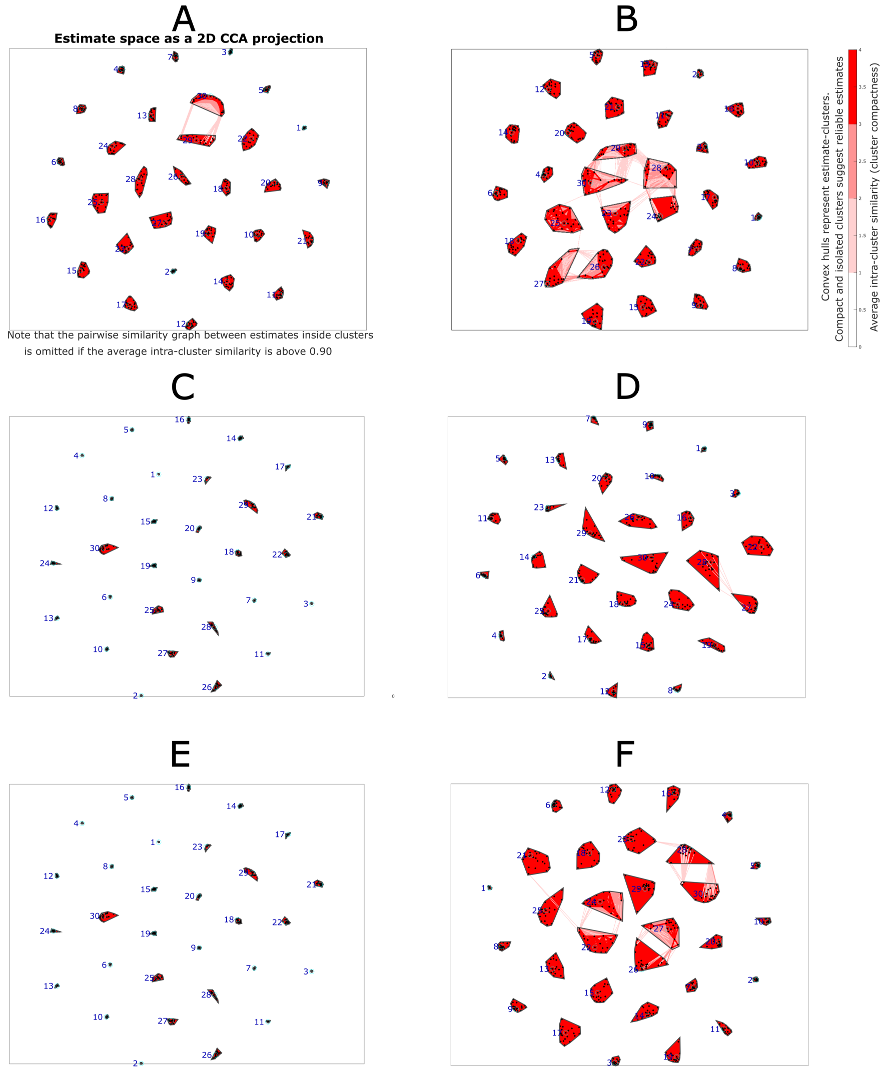


Figure S3: ICASSO visualization of defined clusters as 2D CCA projection for: **(A)** whole data set, electrode-space, **(B)** whole data set, source-space, **(C)** subset 1, electrode-space, **(D)** subset 1, source-space, **(E)** subset 2, electrode-space, **(F)** subset 2, source-space.

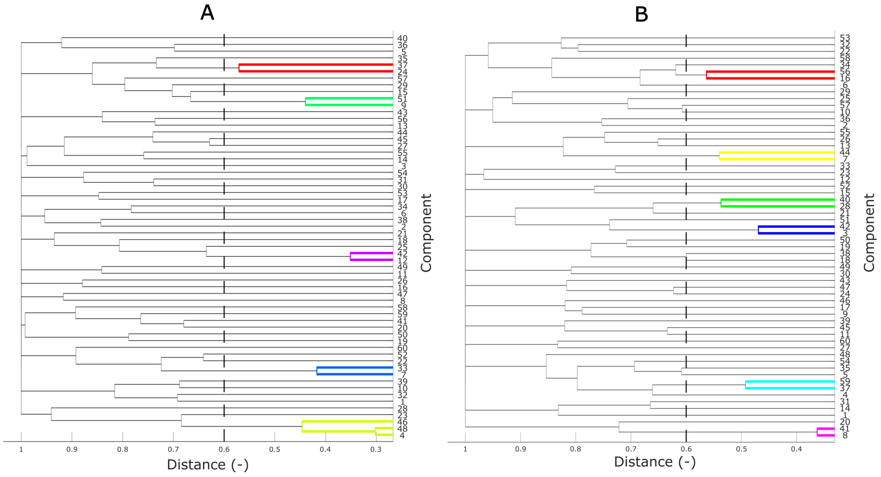


Figure S4: Hierarchical clustering dendrogram of components from both subsets labeled as 1-30 for components from the first subset and 31-60 as components from the second subset with nodes definition by a shortest distance method. The distance metric between components is defined by Equation 4 in the main text. Dashed line denotes a distance cut-off to define clusters both in **(A)** electrode-space, and **(B)** source-space spatio-spectral decompositon.
